# GECSI: Large-scale chromatin state imputation from gene expression

**DOI:** 10.1101/2025.10.02.679828

**Authors:** Jingyuan Fu, Jason Ernst

## Abstract

Compendiums of chromatin state annotations based on integrating maps of multiple epigenetic marks such as from ChromHMM have become a powerful resource. While these compendiums have coverage of many biological samples, there are many additional biological samples that have gene expression data but lack epigenetic mark data and chromatin state annotations. The EpiAtlas resource of the International Human Epigenome Consortium (IHEC) contains a large compendium of chromatin state annotations for which many samples have matched gene expression data, which provides the opportunity to use it to train models to predict chromatin state annotations in additional biological samples with only gene expression data available. To address this, we develop Gene Expression-based Chromatin State Imputation (GECSI), which uses a multi-class logistic regression model trained using a large compendium of gene expression and chromatin state annotations, and apply it to IHEC data. Using cross-validation, we find that GECSI accurately predicts chromatin state assignments and generates probability estimates that are predictive of observed chromatin states, overall outperforming multiple other alternative and baseline methods. GECSI-predicted chromatin states reflect relationships among biological samples and show similar transcription factor and gene annotation enrichments as observed chromatin states. Using available IHEC gene expression data, we apply GECSI to predict chromatin state annotations for 449 additional epigenomes. We expect these predicted annotations and the GECSI software will be a useful resource for chromatin state analyses in many additional biological samples.

## Introduction

Chromatin state modeling based on integrating information from multiple different epigenetics marks, such as histone modifications, through methods such as ChromHMM^1^ is a widely used approach for annotating the genome in a cell type or tissue-specific manner^2^. Chromatin states can correspond to different classes of putative enhancer, promoter, transcribed, repressed, and other types of genomic regions. Annotations of chromatin states have been used for studying gene regulation^3,4,5^, cell and tissue development^6,7^, and phenotype-associated genetic variants^1,8,9,10,11,12^, among other applications.

Multiple compendia of chromatin state annotations have emerged, defined for increasing numbers of reference epigenomes^2,13,14^. Notably, the International Human Epigenome Consortium (IHEC) has recently produced chromatin state annotations for about 1700 reference epigenomes based on six histone modifications using ChromHMM as part of its EpiAtlas resource. In cases where a reference epigenome did not have observed data available for all six histone modifications, but had at least one histone modification experimentally mapped, computationally imputed ChIP-seq data of individual histone modifications were used for the missing experiments. The imputed histone modification data was generated by applying ChromImpute^15^ to the IHEC compendia of ChIP-seq data. Among the samples with chromatin states defined, over 400 of them have both RNA-seq data and chromatin state annotations defined directly based on observed ChIP-seq data for all six histone modifications. The availability of a large compendium of matched chromatin state annotations and gene expression data presents the opportunity to develop and apply an approach to use gene expression data to accurately impute chromatin state annotations for additional biological samples.

While existing chromatin state annotations provide substantial coverage of biological samples, there still exist many samples that have gene expression data available but not chromatin state annotations. Input sample requirements or budget constraints may lead to gene expression profiling but not ChIP-seq assays being conducted in a sample. Within the IHEC compendia, more than 400 biological samples have gene expression data available, but no ChIP-seq data. In addition, many more biological samples have expression data available in public repositories such as the Gene Expression Omnibus (GEO)^16^ without data from ChIP-seq or other epigenetic assays. The ability to effectively impute chromatin state annotations from gene expression thus could greatly expand the number of samples with chromatin state annotations available.

A number of methods have been developed, primarily focusing on the problem of imputing individual epigenomic tracks when tracks for other epigenomic marks are available in a sample, including ChromImpute^15^ and others^17–19^. Methods have also been developed to utilize gene expression data to predict data from assays that profile chromatin accessibility, specifically ATAC-seq or DNase-seq^20–22^. In addition, methods have been developed to learn the relationship between gene expression and histone modifications or other chromatin features^23–26^, mainly focusing on predicting gene expression from epigenetic signals. However, imputing chromatin state annotations from gene expression data, or even individual histone modifications has not been a focus of prior work.

To address this, we introduce Gene Expression-based Chromatin State Imputation (GECSI), a computational method to impute chromatin state annotations based on only gene expression data leveraging the large set of IHEC samples with chromatin state annotations and matched gene expression data. To do this, GECSI trains multi-class logistic regression models. The features of the classifiers are based on the chromatin state annotations in the most similar samples to the target sample as determined by gene expression information. Then, given any new sample with gene expression data, GECSI predicts chromatin state annotations for the sample. By comparing GECSI’s predictions to experimentally derived chromatin state annotations in the IHEC compendia, we observe that overall GECSI makes accurate predictions and compares favorably to several baseline methods or alternative approaches. We expect the additional chromatin state annotations we have predicted in the IHEC epigenomes along with future applications of GECSI will facilitate chromatin state analyses in many additional biological samples.

## Results

### Overview of GECSI

GECSI predicts genome-wide chromatin state annotations for a given sample based on its gene expression data. GECSI generates both probabilistic (“soft”) and discrete (“hard”) annotations. To accomplish this GECSI trains an ensemble of models using a reference compendium containing matched gene expression and chromatin state data **(Figure 1A)**. Specifically, GECSI trains a multi-class logistic regression model for each sample in the compendium that has both gene expression and chromatin state annotation. A sample for which a model is trained on is defined here as a “training sample.” During the training process, GECSI first identifies the *L* nearest samples to a training sample based on gene expression correlation (Methods). GECSI then defines a set of features corresponding to the chromatin state annotations for the ranked position of the neighbor (e.g. first neighbor, the second nearest neighbor, etc) up to the *L*th nearest neighbor. The chromatin state annotations of these neighbors are represented with a binary feature for each combination of neighbor and state, corresponding to the presence or absence of the chromatin state annotation at the genomic position in the neighbor. Next, using these features GECSI trains the multi-class logistic regression models. Such models are trained for all reference samples, and can then be applied for predictions in new samples **(Figure 1B)**. For predictions in a new sample, GECSI finds its *R* nearest neighbors in the reference compendium based on gene expression correlation, and then makes predictions using the *R* corresponding previously trained models. Each model makes probabilistic predictions of the chromatin state at each position. To generate probabilistic chromatin state annotations for new samples, GECSI averages the probabilistic predictions from the individual models. GECSI then determines the hard chromatin state assignment for each genomic position by selecting the state with the highest average probability **(Figure 1C)**.

**Figure 1:**
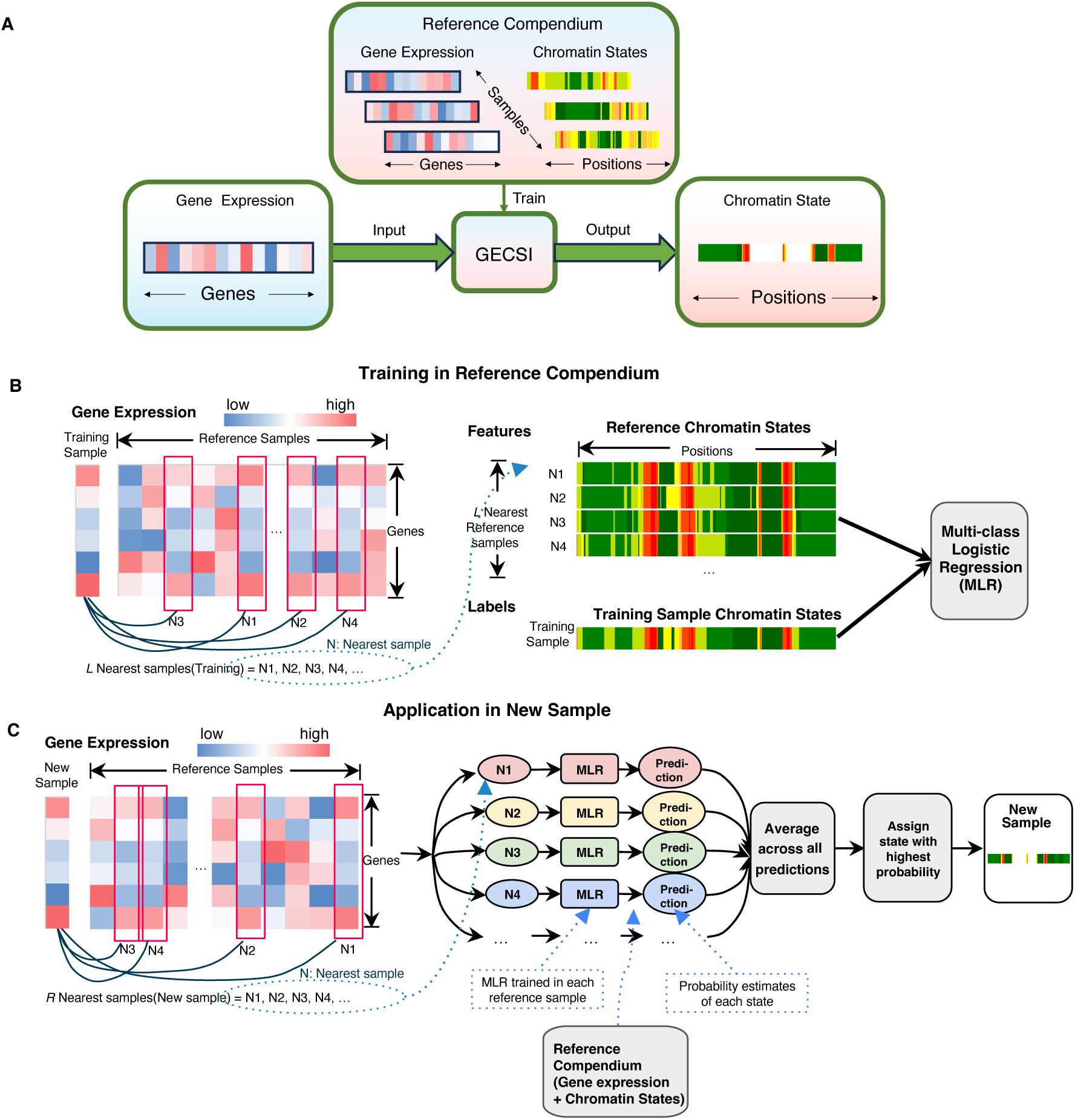
Overview of GECSI. (A) Workflow overview: GECSI is trained using a reference compendium containing both chromatin state tracks and gene expression data. Once trained, GECSI can predict chromatin states for any new sample using its gene expression profile. **(B) Training in reference compendium:** GECSI trains an ensemble of logistic regression models, with each model trained on one sample in the reference compendium, with features generated from a subset of *L* nearest reference samples. For each logistic regression model, one reference sample is designated as the training sample. The model is trained to predict chromatin state annotations for the training sample based on chromatin state and gene expression data from the remaining reference samples, along with gene expression data in the training sample. For a given training sample, GECSI ranks other samples based on gene expression correlation across samples in the compendium. GECSI then extracts chromatin state features based on the samples with the highest correlation. **(C) Application in a new sample:** For a new sample, GECSI selects the *R* nearest reference samples based on gene expression similarity. GECSI then averages the probabilistic predictions of the individual multinomial logistic regression (MLR) models previously trained on the reference samples. Finally, GECSI generates a chromatin state track of hard assignment of states, which is based on the state with the highest probability for each genomic position.

### Visualization of GECSI-predicted chromatin state tracks for IHEC epigenomes

We first applied GECSI using the subset of reference epigenomes that are part of the IHEC compendium and have experimentally mapped ChIP-seq data and gene expression. Specifically, we first focused on training and evaluating GECSI using as samples a subset of 414 reference epigenomes that had experimentally mapped ChIP-seq data for six histone modifications (H3K27ac, H3K27me3, H3K36me3, H3K4me1, H3K4me3, and H3K9me3) along with gene expression data, which we refer to as fully observed epigenomes (**fullobs**). For these epigenomes we used chromatin state annotations based on experimentally mapped ChIP-seq data for the six histone modifications, which we refer to as observed chromatin state annotations. These chromatin state annotations were based on a previously generated 18-state ChromHMM model^2^ and defined at 200-bp resolution. Using GECSI, we made predictions for all the fullobs epigenomes and compared the predictions to observed chromatin state annotations in 5-fold cross-validation. We first confirmed that the fraction of the genome covered by each state is highly correlated with the observed chromatin states (Pearson 0.974). We note that some of the smaller states did not receive a predicted hard assignment annotation in every epigenome, though in all epigenomes they were still detectable in aggregate genomewide based on the soft assignments **(Supplementary Figure 1)**.

To further examine GECSI’s cross-validation predictions, we visually compared GECSI-predicted chromatin state annotations with the corresponding observed chromatin state annotations at randomly sampled 200 kb regions **(Figure 2)**. We observed that at these loci GECSI identifies both constitutive and cell-type-specific chromatin state patterns matching well with ChromHMM-generated chromatin state tracks based on observed data. We have also made available genome browser tracks of GECSI-predicted chromatin state for viewing other genomic loci (see Data Availability).

**Figure 2:**
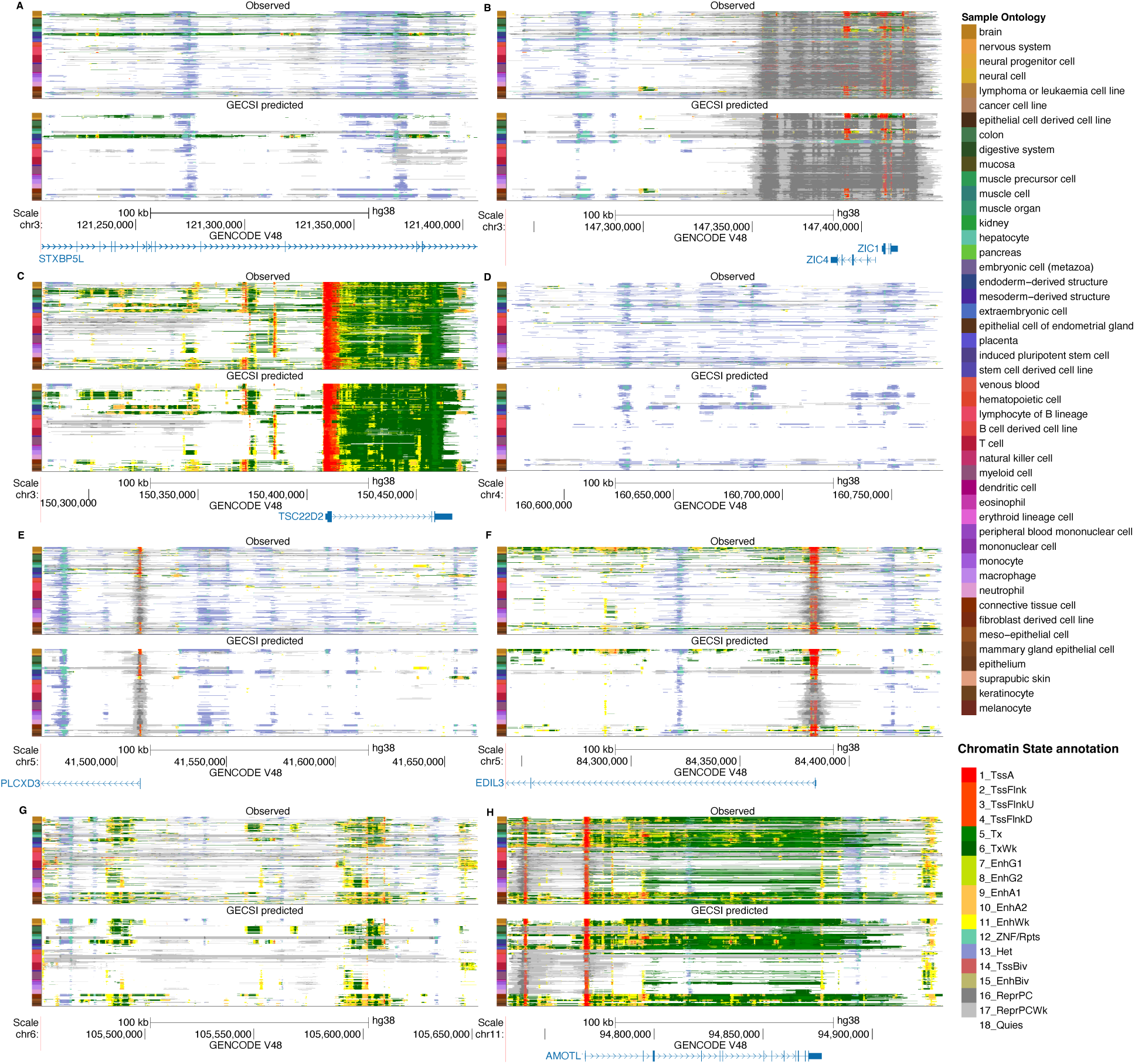
Visualizations of GECSI-predicted and observed chromatin state annotations in randomly sampled regions. Visualization of chromatin state tracks of 414 fully observed IHEC reference epigenomes based on observed data (top) and GECSI’s predictions (bottom) for eight randomly sampled 200 kb regions: **(A)** chr3: 121206801 - 121406800, **(B)** chr3: 147237201 - 147437200, **(C)** chr3: 150278201 - 150478200, **(D)** chr4: 160573401 - 160773400, **(E)** chr5: 41465201 - 41665200, **(F)** chr5: 84242801 - 84442800, **(G)** chr6: 105452601 - 105652600, **(H)** chr11: 94732201 - 94932200. Each row is a chromatin state track for an IHEC reference epigenome. Tracks are ordered by a previously defined ordering by IHEC. The color label on the left is based on intermediate sample ontology groups from the IHEC metadata. Annotations generated by UCSC genome browser with GENCODE v48 and Genome assembly GRCh38. Chromatin state annotations follow the color coding as previously2 and the color coding is shown in the lower right corner, with detailed descriptions of the annotations provided in the supplement **(Supplementary Table 1)**.

### Agreement of GECSI’s chromatin state predictions with observed chromatin state annotations

We next quantitatively evaluated GECSI’s chromatin state predictions with that of the observed chromatin state annotations. For these evaluations, we developed or used several alternative and baseline methods for comparison. As a simple baseline method we compared to the **Majority state** baseline, which selects the most frequent state across all samples at each genomic position. We also developed and compared against two other methods, which also had not been used for this problem. The first is the **k-nearest-neighbor (KNN)** method, which assigns the most frequent state across the *k* nearest samples as determined based on gene expression correlation (Methods). The other method which we term **ChromImpute+ChromHMM** is based on applying ChromImpute^15^ followed by ChromHMM^1^. For this we designed an approach to use ChromImpute to impute ChIP-seq data using only gene expression information from the target sample along with the reference compendium (Methods). We then binarized these imputed histone modification data, which we used as input to ChromHMM to generate both soft and hard chromatin state assignments based on the same 18-state model (Methods). We evaluated GECSI and the other methods using cross-validation.

We first evaluated how predictive the soft assignments of GECSI are of observed chromatin state hard assignments and compared it to the alternative methods and baselines. For KNN we used the frequency of the state in the K-nearest epigenomes in the comparisons and for the Majority State the frequency of the state at the position overall. We also compared GECSI’s soft assignments with the posterior probability assignments as computed by ChromImpute+ChromHMM (Methods). For these comparisons, we used Area-Under the Precision-Recall Curve (AUPRC) and Area-Under Receiver Operating Characteristic curve (AUROC) metrics when predicting the observed chromatin state hard assignments. We computed the AUPRC and AUROC values separately for each combination of 414 fully observed epigenomes and 18 chromatin states, giving 7,452 comparisons in total **(Figure 3)**. Among all combinations, GECSI achieved a higher AUPRC and AUROC than the alternative methods evaluated. GECSI was better than the next best method, KNN for AUPRC and Majority State in AUROC, for 82.6% and 79.0% of all combinations **(Figure 3D)**, respectively. The overall distribution of scores from GECSI were also significantly better than that from the second best method in each evaluation (p-value of < 2 × 10^-16^, based on a one-sided Wilcoxon signed-rank test; effect size of improvement based on Cohen’s d of 0.134 and 0.314 for AUPRC and AUROC, respectively). We also evaluated on a per-state basis whether GECSI had a better overall distribution of scores and observed that GECSI achieved statistically significant improvement over all other alternative methods for 16 of the 18 chromatin states based on AUPRC and 15 of the 18 chromatin states based on AUROC **(Figure 3B, Supplementary Figure 2-3)**. These results support that overall the rankings from GECSI’s soft assignments are more predictive of observed chromatin state hard assignments than those based on other methods considered.

**Figure 3:**
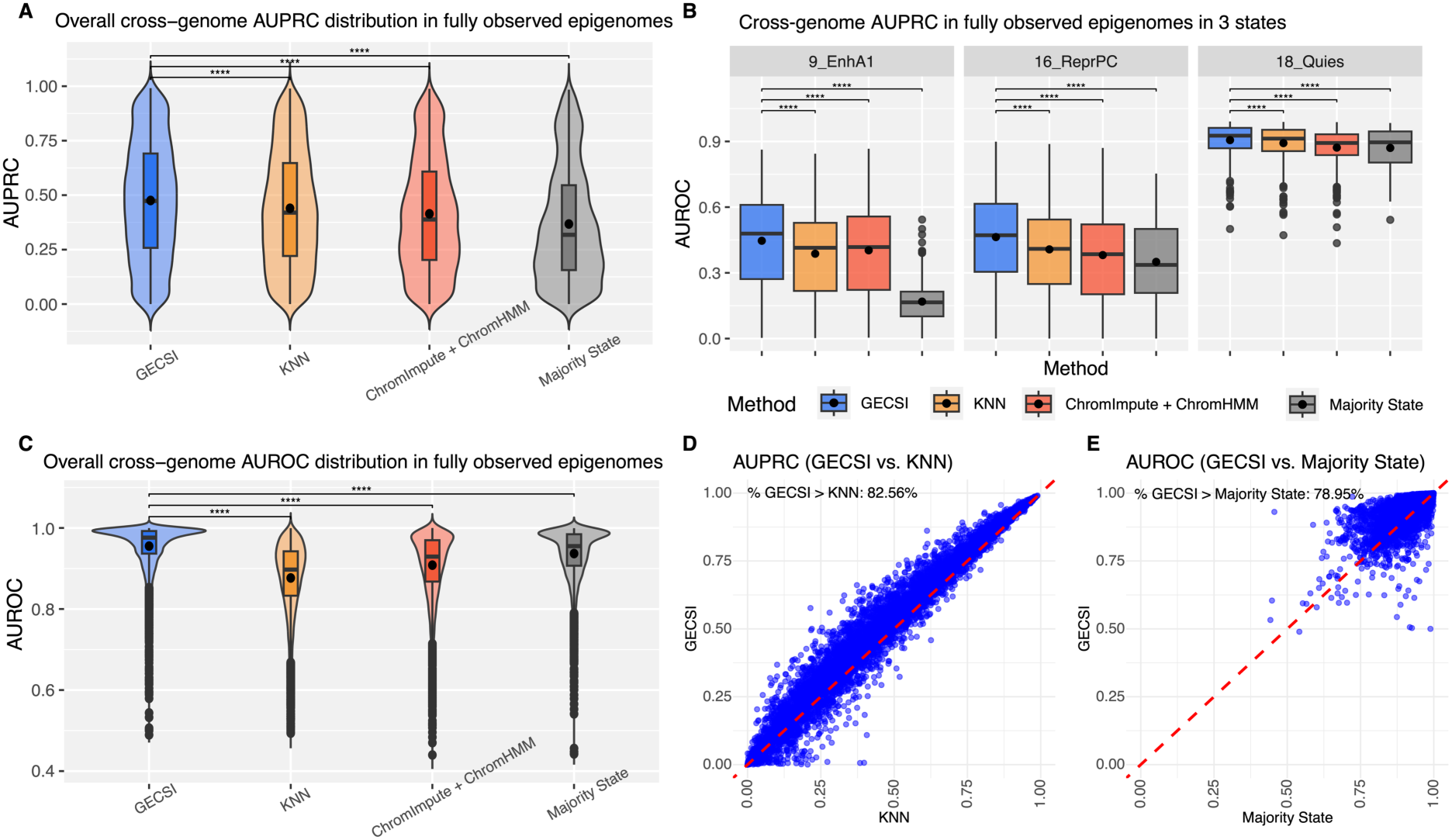
**Evaluations of chromatin state soft assignment predictions in the fullobs epigenomes**. Shown are cross-validation results for soft assignments predicting 18 chromatin states among 414 fullobs reference epigenomes, specifically: **(A)** Violin and boxplots showing the distribution of AUPRC across all combinations of epigenomes and chromatin states separately for GECSI, KNN, ChromImpute+ChromHMM, and Majority State. **(B)** Boxplots displaying AUPRC distribution for three chromatin states, Active Enhancer 1 (9_EnhA1), Repressed Polycomb (16_ReprPC), and Quiescent/Low (18_Quies) for each method evaluated. Results for additional states can be found in **Supplementary Figure 2. (C)** Violin and boxplots showing the distribution of AUROC across all combinations of epigenomes and chromatin states separately for GECSI, KNN, ChromImpute+ChromHMM, and Majority State. For (A) to (C), statistical significance of difference between GECSI and other methods was assessed using one-sided Wilcoxon signed-rank test, and significant differences are indicated with asterisks (*: p < 0.05, **: p < 0.01, ***: p < 0.001, ****: p < 0.0001). Each box represents the interquartile range (IQR) from the 25th to 75th percentile, with the horizontal line inside the box indicating the median, and the dot inside the box indicating the mean. Whiskers extend to the smallest and largest values within 1.5 × IQR from the lower and upper quartiles, respectively; points beyond this range are shown as outliers. **(D)** Scatterplot comparing AUPRC scores of GECSI (y-axis) and the method with the second best AUPRC, KNN, (x-axis) with the red dashed diagonal line representing y=x. The percentage of points with higher GECSI compared to KNN is represented in the upper left corner of the plot. **(E)** Scatterplot comparing AUROC scores of GECSI method (y-axis) and the method with the second best AUROC, Majority State (x-axis), with the red dashed diagonal line representing y=x. The percentage of points with higher GECSI compared to Majority State is represented in the upper left corner of the plot.

We also evaluated and compared the performance of GECSI’s hard assignment predictions (Methods). One metric we used for the evaluations was the overall accuracy. We also used the weighted F1 score and weighted Jaccard index which were weighted averages of the F1 score and Jaccard index, respectively, computed on a per-state basis with weights determined by state size (Methods). GECSI obtained an average accuracy of 0.726 and weighted Jaccard and F1 averages of 0.586 and 0.706, respectively. In comparison, GECSI’s overall accuracy was significantly better than the best-performing alternative or baseline method, KNN, which obtained an average accuracy of 0.716 (p-value < 8 ×10^-15^, effect size 0.11). For weighted Jaccard and F1 averages, GECSI’s performance was comparable to the best-performing alternative or baseline method, also KNN for both, which had a weighted Jaccard and F1 average of 0.582 and 0.708, respectively **(Supplementary Figure 4-6)**. To better contextualize the hard assignment evaluation performance we also compared GECSI with a “closest reference” benchmark. The “closest reference” is defined as the reference epigenome that has the highest proportion of shared chromatin state assignments (Methods). We note the “closest reference” benchmark is not an actual method since it requires knowledge of the target chromatin state annotations that are being predicted. Overall, the predictive performance of GECSI in these evaluations closely approached that of the closest reference benchmark, which had values of 0.734, 0.602, and 0.725 for accuracy, weighted average Jaccard, and weighted F1, respectively. These results support the overall effective performance of GECSI for chromatin state prediction.

Although often accurate, GECSI’s predictions still did not match observed annotations for a substantial fraction of the genome. We hypothesized that in some cases this could be due to GECSI predicting a similar but not the same chromatin state as the observed chromatin state annotations. To investigate this, we first manually defined groups of chromatin states based on similar biological interpretation (Methods). We observed that among all positions where the observed chromatin state annotation did not match the predicted, on average 18% of the time the observed state was still within the same group as the predicted state, which was 8.8 fold higher than expected by chance (Methods; **Supplementary Figure 7**). When further restricted to positions that had non-quiescent GECSI predictions or restricted to non-quiescent annotations based on observed data, the corresponding percentages increased to 38% and 23%, respectively, which were 2.9 and 3.8 fold higher than expected by chance. These results suggest that for positions where GECSI’s prediction does not match the observed chromatin state annotation, GECSI will still in many cases predict a similar chromatin state assignment.

We also evaluated the extent to which GECSI’s probabilistic predictions are well calibrated. To do this we grouped genomic positions based on having a similar probabilistic prediction value for the presence of a specific chromatin state and then evaluated the extent to which the prediction value agreed with the observed proportion of the state among those positions (Methods). We observed strong correlations between predicted probabilities and observed fractions (average Pearson correlation of 0.91 when averaged across all epigenomes and chromatin states) (**Figure 4A, Supplementary Figure 8**). Although overall well calibrated, we did observe places where GECSI’s predictions were under confident, particularly for smaller chromatin states. In addition to overall calibration, we investigated if GECSI’s probabilistic predictions were informative about the presence of specific observed states other than the one being predicted (**Figure 4B, Supplementary Figure 9**). Specifically for cases in which a GECSI-predicted state was not observed, we evaluated the extent to which GECSI-predicted probabilities of a state being present correlated with a similar state being observed. For this we used the same state group definitions as used above to define similar states and found a strong correlation on average across chromatin states (Spearman mean 0.63).

**Figure 4:**
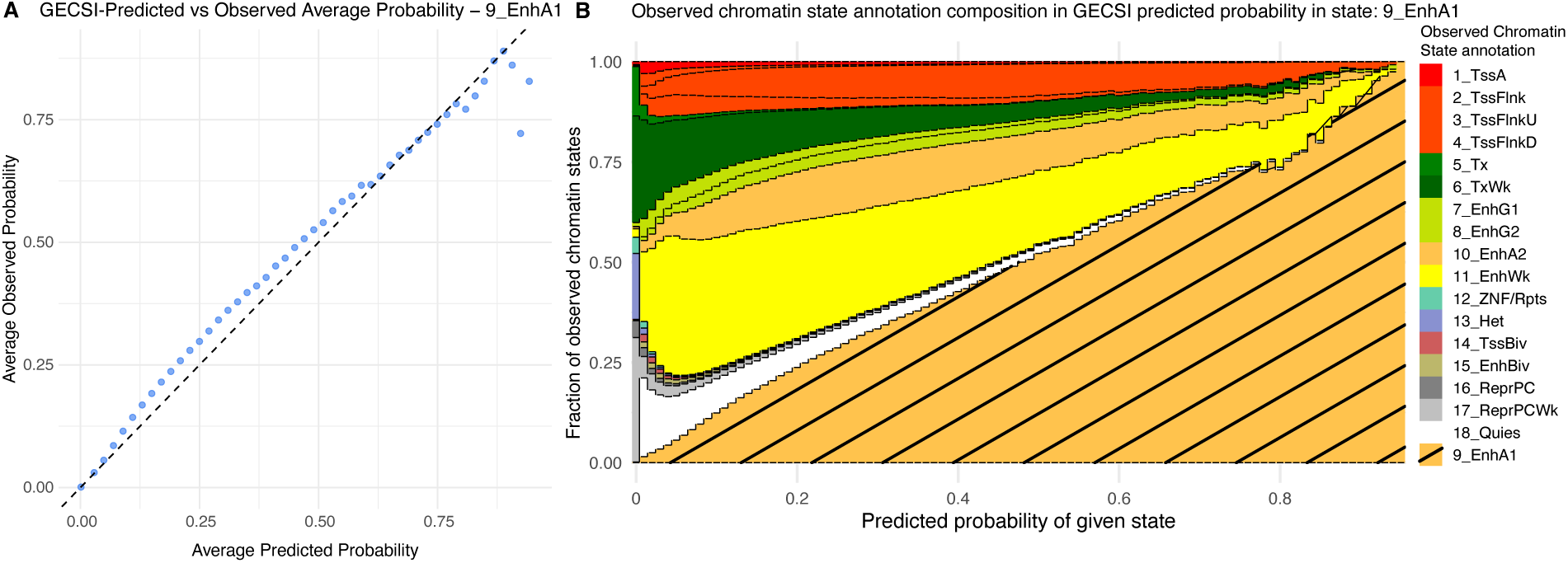
**Calibration analysis of GECSI-predicted probabilities**. **(A)** Scatterplot comparing average GECSI-predicted probabilities for the Active Enhancer 1 (9_EnhA1) state (x-axis) to the average observed frequency of the state (y-axis) from a cross-validation experiment among 414 fully observed reference epigenomes. Genomic positions were grouped into 50 equally spaced bins based on probability predictions. Average predicted probabilities and average observed probabilities are computed across all positions in each bin and all epigenomes. The diagonal black dashed line marks the y=x line, representing perfect calibration. Similar plots for other chromatin states are available in **Supplementary Figure 8. (B)** Observed chromatin state annotation composition with respect to predicted probabilities of the given state, Active Enhancer 1 (9_EnhA1), which are grouped into 100 bins based on the predicted probability of the given state shown on the x-axis. Fraction bars are colored by the color-coding of chromatin states. The color bar corresponding to state 9_EnhA1 is located at the bottom of each bar with diagonal shading overlaid for better visualization, while the other states are stacked according to the order of the chromatin states. Similar plots for other chromatin states are available in **Supplementary Figure 9**.

### Enrichments of GECSI-predicted chromatin states for genome annotations and TF binding

We next evaluated the consistency of the enrichments of GECSI-predicted and observed chromatin state annotations for a selected set of external cell type invariant genome annotations. The genome annotations included transcription start sites (TSS), gene bodies, exons, transcription end sites (TES), regions within 2kb of a TSS, and CpG islands. The enrichments based on the two chromatin state annotation sets were overall highly concordant. For instance when considering TSS annotations, we observed the greatest fold enrichment in the Active TSS state (1_TssA) based on GECSI’s predicted chromatin state annotations and inferred based on observed data, 70.7 and 71.0 fold, respectively. When considering CpG islands, we found the greatest enrichment based on both sets of annotations for the Bivalent Promoter state (14_TssBiv) state, which had fold enrichments of 102 and 88.1 respectively for chromatin state annotations based on GECSI’s predictions and the observed data, respectively (**Supplementary Figure 10**). For each of the six external categories that we considered, we computed for each cell type the correlation of enrichments for GECSI-predicted chromatin state annotations and those inferred directly based on observed data and then averaged those correlation values. On average across the external categories, we obtained an average Pearson correlation of 0.945 between enrichment scores computed in observed chromatin states and those computed in GECSI-predicted chromatin states, with the highest correlation being 0.965 for gene body annotations and the lowest correlation 0.914 for exons annotation. These results suggest that enrichment analyses for cell-type invariant features using GECSI-predicted chromatin states will yield similar results to using observed data.

We also investigated the extent to which enrichments based on GECSI’s predicted chromatin state annotations are consistent with those based on observed data for a class of cell type variant features, specifically locations of transcription factor (TF) binding. For this we conducted enrichment analyses in six different cell lines with a total of 1500 TF binding experiments collectively obtained across those same cell lines (Methods). We found that GECSI-predicted chromatin state annotations exhibited different enrichments for TF binding that were largely consistent with enrichments based on observed chromatin state annotations **(Figure 5)**. Overall, the Pearson correlation between enrichments based on the observed and predicted chromatin state annotations was on average 0.84 across the cell lines **(Methods; Supplementary Figure 11)**. We also confirmed consistent with what has been previously established^27,28^, that when using GECSI-predicted chromatin state annotations, TF binding enrichments for promoter states are still more constitutive and less dynamic across cell types compared to enhancer states. For example, the Active TSS state (1_TssA) showed strong enrichment for TF binding regardless of whether the TF binding experiment was in the same cell type of the chromatin state annotations or a different one, with median fold enrichments of 61.4 and 59.9, respectively. These high fold enrichments are consistent with the result found based on observed chromatin state annotations, which were 51.7 and 48.8, respectively. Compared to promoter chromatin state annotations, GECSI’s predicted enhancer chromatin state annotations showed greater differences in enrichments when the TF binding experiment was conducted in the same cell type as the chromatin state annotations than a different one. For example, for Active Enhancer state (9_EnhA1) the median fold enrichments for TF binding experiments were 23.2 and 8.4 fold when conducted in the same or different cell type as the chromatin state annotations, respectively. This difference in enrichment scores is also consistent with differences based on observed chromatin states, 22.3 and 5.3 fold, respectively (**Supplementary Figure 12A**). We observed similar overall enrichment patterns when conducting the analysis using soft-assignments after truncating low probabilities to zero (**Methods; Supplementary Figure 12B**). These results suggest that enrichment analysis for cell-type variant features using GECSI-predicted chromatin state annotations also yields similar results to using chromatin state annotations based on observed data.

**Figure 5:**
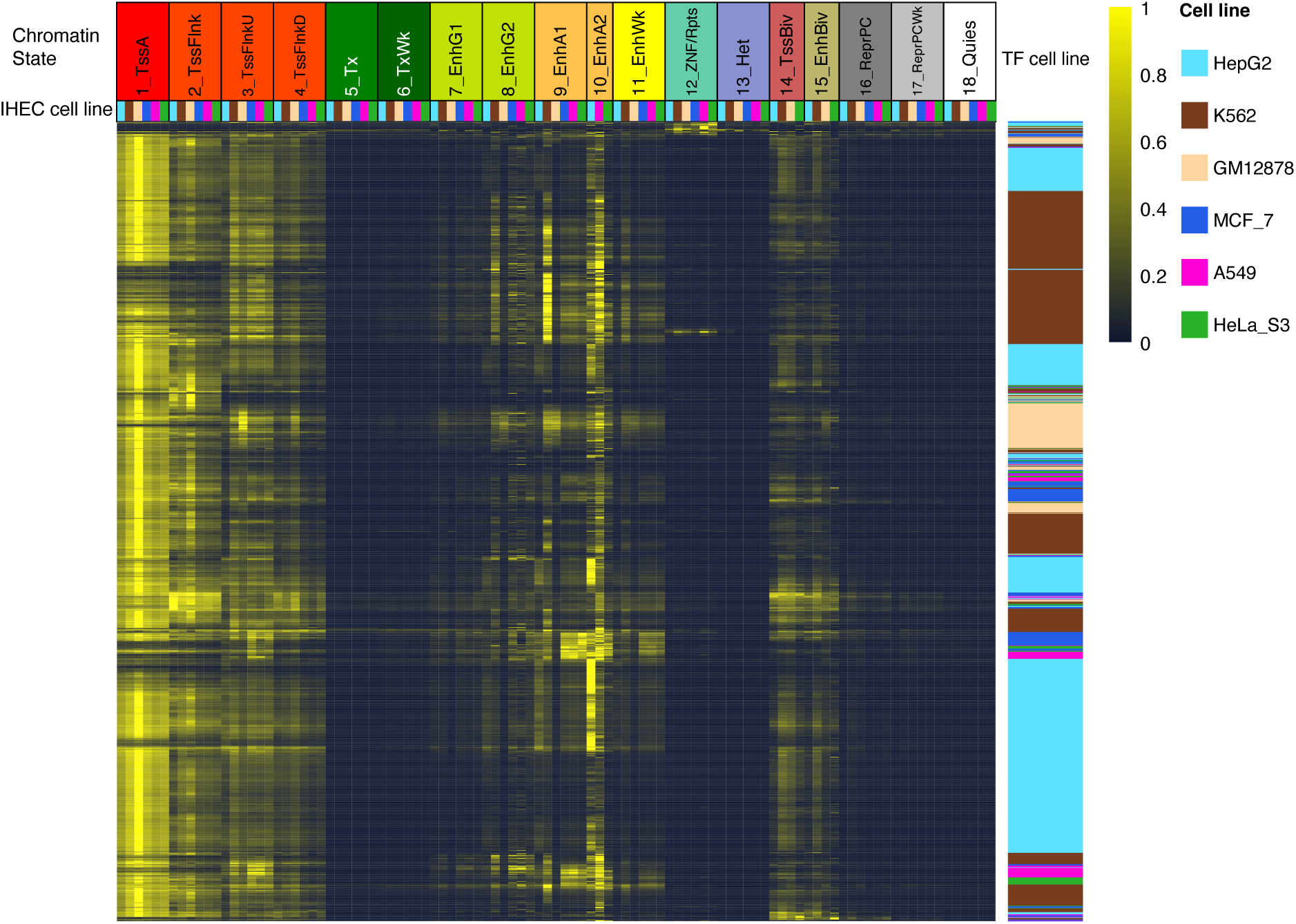
Transcription factor (TF) binding enrichments in different GECSI-predicted chromatin states across six cell lines. TF binding enrichment heatmap: Each row corresponds to a TF ChIP-seq experiment in a given cell line. Each column corresponds to a chromatin state-cell line combination. The entries in the heatmap are fold enrichments normalized between 0 and 1 for each row separately, where the maximum is scaled to 1 and the minimum to 0. The fold enrichments are calculated using GECSI-predicted chromatin state hard assignments and peak calls from TF ChIP-seq experiments. The top right color scale shows the scaled enrichments with yellow representing higher enrichments and blue representing lower enrichments. The ordering of the rows was determined based on the TF binding enrichments in observed chromatin state annotations **(Supplementary Figure 12)** (Methods). The columns are ordered first by chromatin states and then by individual cell lines using the following order: HepG2, K562, GM12878, MCF-7, A549, and HeLa-S3, restricted to those cell lines for which the state is available. The top row of labels correspond to chromatin states and the second row of color labels represent cell lines. The rows are color-coded by the cell line of the TF ChIP-seq experiment, with the same color-coding as the column labels.

### GECSI-predicted chromatin state annotations capture expected epigenome relationships

We also evaluated the extent to which GECSI’s chromatin state predictions capture expected relationships between epigenomes based on observed data. For this we first focused on evaluating sample similarity based on GECSI’s predicted chromatin state annotations and comparing it with sample similarity based on chromatin state annotations from observed data. Specifically, we calculated the Pearson correlation defined on a per-state basis, between epigenomes based on a binary vector representation of hard assignments for both predicted and observed chromatin states (Methods). We saw on average the correlation values were higher based on GECSI’s predicted chromatin state annotations than based on observed chromatin states (0.27 vs. 0.20 on average). While higher, the pairwise sample-wise correlation values based on GECSI’s predictions were still well correlated with those based on observed chromatin state annotations with a Pearson correlation that was as high as 0.77 for an active enhancer state (9_EnhA1) and 0.60 on average across states (**Figure 6A-B; Supplementary Figure 13-15**). We also evaluated the correlations using soft assignment predictions on a per state basis among epigenomes and observed that these correlation values were well correlated with those from observed data (Pearson 0.52) **(Supplementary Figure 16)**. In addition to these evaluations on a per state basis, we also conducted evaluations based on the overall pairwise agreement, which we defined as the overall proportion of positions for which two sets of chromatin state annotations matched. We saw on average higher overall pairwise agreement among samples based on GECSI’s predicted annotations compared to those based on observed data (0.67 vs. 0.54) while still seeing a strong average correlation among the overall pairwise agreement values based on observed and predicted annotations (Pearson correlation 0.70).

**Figure 6:**
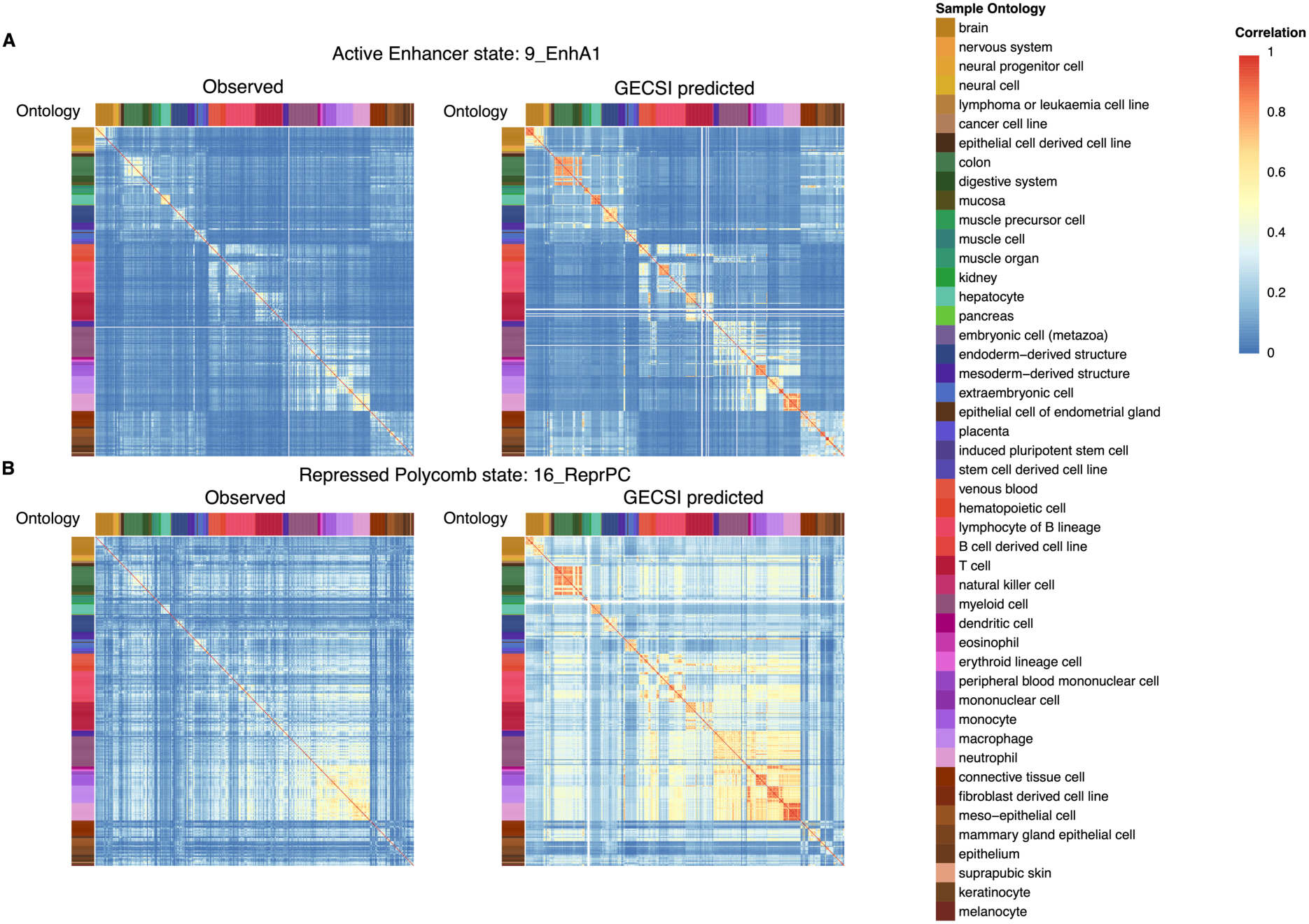
Pairwise sample relationships generated from predicted chromatin states and observed chromatin states. Heatmaps showing pairwise correlation among 414 fully observed IHEC epigenomes in two chromatin states: **(A)** Active enhancer state 1 (9_EnhA1) and **(B)** Repressed Polycomb state (16_ReprPC). In both (A) and (B), left panel shows correlations from observed chromatin state annotations and the right panel shows correlations from GECSI-predicted. Corresponding heatmaps of all other chromatin states are available in **Supplementary Figures 14-15**. Each row and column represents an epigenome. Each grid entry represents a Pearson correlation between the epigenome of the row and the epigenome of the column using the binary presence or absence of the corresponding chromatin state, following the color scale shown on the top right. Entries in the grid are shown as white for epigenomes where the corresponding chromatin state is not available. Correlation results are ordered based on the IHEC epigenome ordering, and color annotation of each epigenome is based on the sample intermediate ontology metadata following the IHEC coloring.

We further investigated the extent to which GECSI’s chromatin state predictions reflect expected relationships between epigenomes based on shared metadata. For this we focused on the intermediate level sample ontology grouping in IHEC’s epigenome metadata. We first evaluated the correlation on a per state basis among epigenomes belonging to the same group and those belonging to different groups using the binary vectors of hard assignment predictions. Correlations based on GECSI’s predictions on average had higher within-group sample-wise correlations compared to the observed chromatin state annotations (0.55 vs 0.35, averaged across chromatin states). Furthermore, the difference of within-group sample-wise correlations to those based on epigenomes belonging to different groups was greater for all 18 chromatin states when using GECSI’s predictions compared to those based on observed data (on average 0.30 vs. 0.15). We also compared GECSI’s predicted chromatin state annotations with observed annotations for each sample ontology group individually in terms of the difference between the average correlation between pairs of epigenomes within the same group with correlations between pairs of epigenomes from different groups. We observed that for 30 of 47 sample ontology groups GECSI obtained a larger difference on average (p-value = 0.039, based on a one-sided binomial test). In addition we evaluated the overall pairwise sample-wise agreement among epigenomes **(Supplementary Figure 17)**. We observed higher within-group sample-wise agreements in GECSI-predicted chromatin states compared to observed chromatin state annotations (0.84 vs 0.65). We also observed an increase in the difference between within-group sample-wise agreements to those belonging to different groups when using GECSI’s predictions compared to the observed chromatin state annotations (on average 0.18 vs 0.11). We further evaluated the extent to which whether two samples are of the same group can be predicted based on overall pairwise agreement using the Area Under the Precision-Recall Curve (AUPRC) (Methods). We obtained higher AUPRC using GECSI’s predictions compared to using observed data (0.55 vs 0.30). These results suggest that GECSI’s chromatin state predictions better capture expected relationships between epigenomes based on metadata than the observed chromatin state annotations.

### Chromatin state predictions in samples with only RNA-seq experiments available

We also applied GECSI to predict chromatin state annotations for 449 epigenomes in the IHEC EpiAtlas that did not have any histone mark experimentally profiled but did have gene expression data available to generate a new annotation resource. We refer to these epigenomes as GECSI-predicted chromatin state only epigenomes, or **predCS** epigenomes in short. We evaluated the extent to which the pairwise relationship between GECSI’s chromatin state predictions for these epigenomes reflected expected relationship based on sample ontology as above. We compared these results to the results seen previously based on chromatin state annotations inferred from observed data in the fullobs epigenomes. We confirmed that consistent with the previous results that when restricted to predCS epigenomes, those from the same group had higher average pairwise Pearson correlations using binary assignments than from different groups (0.62 vs. 0.25) and likewise for pairwise agreement (0.87 vs. 0.70) (**Figure 7A-B, Supplementary Figure 18**). We found similar patterns in pairwise correlations between predCS epigenomes using soft assignments **(Supplementary Figure 19)**. We also correlated GECSI’s hard assignment predictions in the predCS epigenomes with the observed annotations in the fullobs epigenomes and evaluated the extent to which their correlations are higher if they belong to the same group. We observed that GECSI’s predictions in the predCS epigenomes showed higher average Pearson correlation (0.42) and higher sample pairwise agreement (0.73) with the fullobs epigenomes from the same group than those from different groups (0.22 and 0.60 respectively). These results support that for epigenomes which only have GECSI’s predicted chromatin state annotations available, they also capture expected biological relationships between epigenomes.

**Figure 7:**
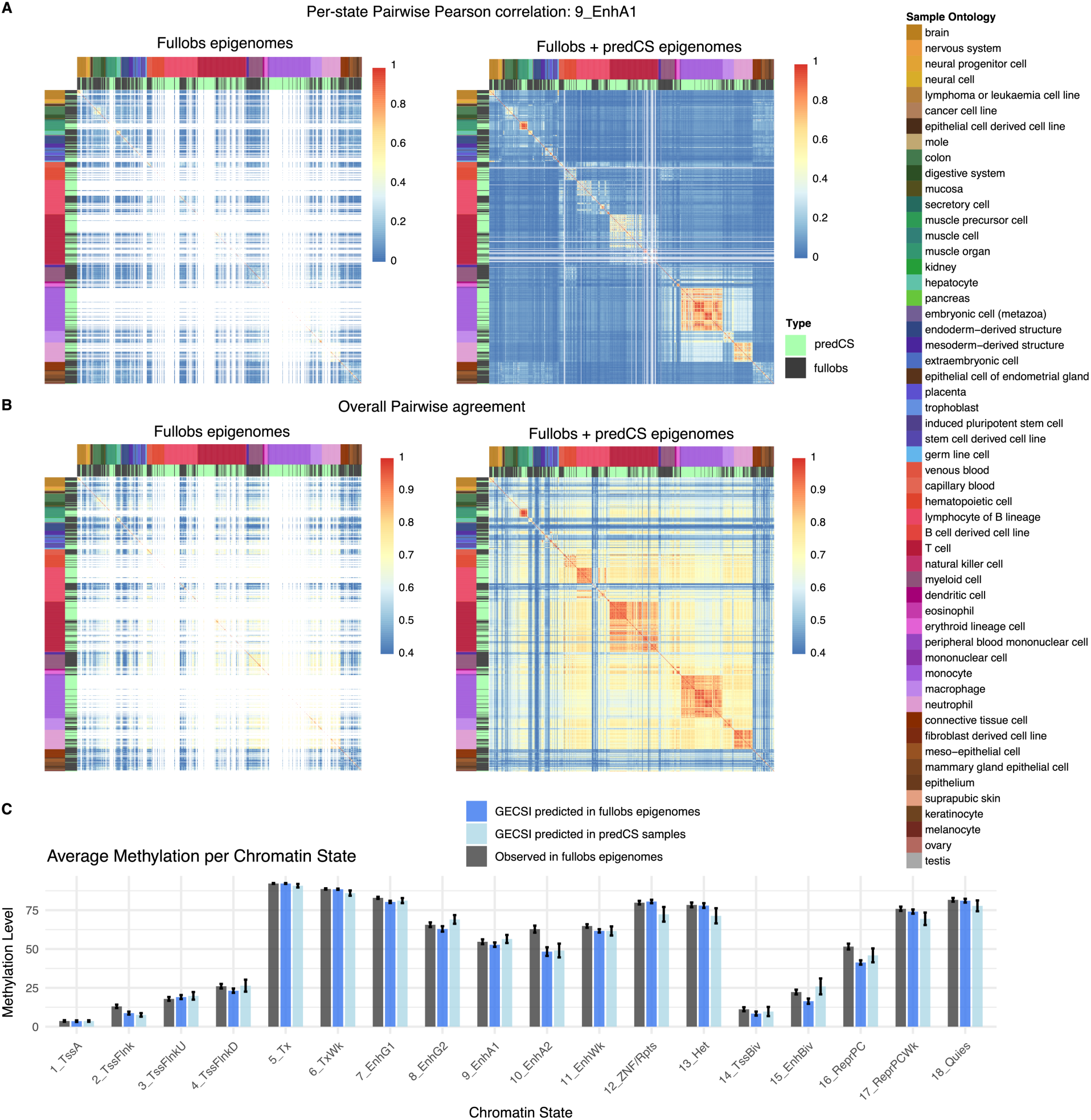
GECSI-predicted ChromHMM annotations only epigenome analysis. **(A)** Pairwise Pearson correlation in Active enhancer state 1 (9_EnhA1) and **(B)** overall pairwise agreement between reference epigenomes generated from observed chromatin states from fullobs epigenomes and GECSI-predicted chromatin states from predCS epigenomes. Each row and column represents an epigenome, and each grid entry represents the corresponding metric calculated between the epigenome on the row and the epigenome on the column using the presence or absence of the corresponding chromatin state, following the color scale on the right side of each panel. The left sub-figure of both (A) and (B) display results based on only observed chromatin states in the fullobs epigenomes, and the right sub-figure of both (A) and (B) display results based on both observed chromatin states in the fullobs epigenomes and imputed chromatin states from predCS epigenomes. In addition, for all heatmaps, epigenomes where the corresponding chromatin state is not assigned are also colored as white. Correlation results are ordered based on the provided sample ordering of IHEC, and color annotation of each sample is based on the sample intermediate ontology metadata (top row) and whether the sample is fullobs (black) or predCS (light green) (second row). The sample ontology color labels are shown on the right side of the full figure. **(C)** Barplot showing the average methylation level for each chromatin state in all chromatin state tracks. Each set of bars represents the average methylation scores for one chromatin state, where the dark gray bars represent observed annotations in the fullobs epigenomes, dark blue bars represent GECSI-predicted annotations in the fullobs epigenomes, and the light blue bars represent GECSI-predicted annotations in the predCS epigenomes’ chromatin state annotations. Bar heights represent the average methylation score across all corresponding epigenomes. The black error bars represent the 95% confidence interval of the average methylation scores.

To further evaluate GECSI’s predicted chromatin state annotations in the predCS epigenomes we focused on DNA methylation data from Whole-Genome Bisulfite Sequencing (WGBS), which was also available in 68 of these epigenomes. Specifically, we sought to establish that the average CpG DNA methylation levels for different chromatin states based on GECSI’s prediction are similar to what is seen based on chromatin state annotations based on observed data, which was indeed the case (**Figure 7C**). This was the case for GECSI’s predictions in both predCS epigenomes (Pearson 0.990) and fullobs epigenomes (0.992). These results demonstrate that GECSI’s predicted chromatin state annotations in both the predCS and fullobs epigenomes reflect DNA methylation patterns seen based on observed chromatin state annotations.

## Discussion

Here we introduced GECSI, a computational method to predict chromatin state annotations from gene expression data using a compendia of samples with both available. Prior approaches to annotate chromatin states in a sample had relied on the availability of other epigenomic data such as ChIP-seq, ATAC-seq or DNase I hypersensitivity experimentally mapped in the sample. Here we applied GECSI in the context of the IHEC EpiAtlas resource to produce predicted ChromHMM annotations for 449 samples that only had gene expression data available, broadening the set of samples which have chromatin state annotations available within the IHEC compendia.

We evaluated GECSI’s soft assignment and hard assignment predictions in cross-validation and compared them to alternative and baseline methods using multiple different evaluation metrics. In most evaluations, GECSI compared favorably. In a subset of evaluations it was the case that GECSI’s performance was similar to the best performing other method compared, but we note that predicting chromatin state annotation directly from gene expression using any of the approaches is novel here. In cases where GECSI’s hard assignments made predictions that disagreed with the observed chromatin states annotation, GECSI would still often predict a similar state assignment. We also showed that GECSI’s soft assignments produced well-calibrated probabilistic predictions that are informative about the presence of specific observed states other than the one being predicted.

We also evaluated whether enrichments obtained using GECSI’s chromatin state annotation predictions are consistent with those based on observed chromatin state annotations. For this we focused on external gene annotations and TF binding and found consistent enrichments in both sets of analyses. This included showing that GECSI’s predictions capture state-specific genomic features including both cell-type-invariant and, importantly, also cell-type-specific features.

Additionally, we evaluated the extent to which GECSI’s predictions capture expected pairwise sample relationships. For this we conducted evaluations based on both the consistency with relationships based on observed data as well as based on sample metadata. We found that GECSI’s predictions generally captured similar pairwise relationships as in the observed data and showed even stronger consistency with sample metadata than the chromatin state annotations based on the observed data.

While we have demonstrated GECSI to be effective, there are limitations and directions for future work. One limitation is that we currently only applied GECSI to bulk gene expression. Given the extensive availability of single-cell gene expression data and limited corresponding chromatin state annotations, a potentially important future direction would be to apply and possibly adapt GECSI to make predictions based on single-cell gene expression data. One challenge with single-cell applications compared to in the bulk setting is evaluating the effectiveness of predictions given the limited chromatin state annotations available based on single-cell data. Another limitation and future direction is that while GECSI is overall accurate, there is potential room for improvement particularly for small chromatin states. We also note that certain states did not receive hard assignments in every epigenome, though were still detectable based on soft-assignments. A direction for future work could be to investigate deriving hard assignments based on sampling from GECSI’s state probability estimates instead of always using the maximum probability assignment. One potential route for improving GECSI’s predictions is to leverage additional data for training in particular epigenomes that did not have observed data for all six histone modifications, but had imputed data for the missing ones. We did not use data from such epigenomes primarily to ensure in our cross-validation evaluation procedures there was no data leakage between training and testing folds as could happen through the use of imputed histone modification data defined based on the full compendium. Another direction for future work would be to apply GECSI in additional species where large datasets of gene expression and matched chromatin state data are available, such as in mouse^29,30^.

While these directions should further enhance the utility of GECSI, we have already demonstrated that GECSI could substantially expand the number of IHEC samples with chromatin state annotations available. We expected these predicted chromatin state annotations combined with future applications of GECSI method will be a resource to conduct chromatin state analyses in a broader range of biological samples.

## Materials and Methods

### IHEC data preprocessing

We obtained processed chromatin state annotations, ChIP-seq data and RNA-seq data from the IHEC EpiAtlas compendium. The ChIP-seq data was for six histone modifications: H3K27ac, H3K27me3, H3K36me3, H3K4me1, H3K4me3, and H3K9me3. For the reference epigenomes that have all six experimentally mapped histone modifications, 414 also have one of the RNA-seq assays: mRNA-seq or total-RNA-seq. These 414 epigenomes are the ones we refer to as fully observed epigenomes, or fullobs epigenomes for short, as their chromatin states come defined based on experimentally mapped data for all six histone modifications. We used these 414 epigenomes for training GECSI and cross-validation evaluations. In addition, we applied GECSI to 449 epigenomes with RNA-seq data available, but no ChIP-seq data available. These are the epigenomes we refer to as GECSI-predicted chromatin state only epigenomes, or predCS epigenomes for short.

We obtained ChromHMM^1^ chromatin states annotations from IHEC for the 414 fullobs epigenomes. The ChromHMM annotations were generated using the 18-state model originally from the Roadmap Epigenomics Consortium^2^. The chromatin state annotation tracks were defined at 200 base pair resolution, which was the same resolution that we used for training and evaluating GECSI.

Among the 414 fullobs epigenomes, 176 epigenomes had only mRNA-seq available, 201 had only total RNA-seq available and 37 epigenomes had both mRNA-seq and total-RNA-seq. Among the 449 predCS epigenomes, 91 epigenomes had only mRNA-seq available, 341 had only total-RNA-seq available, and 17 epigenomes had both mRNA-seq and total-RNA-seq. For those epigenomes with both RNA-seq assays available, we used mRNA-seq. We obtained the Transcripts Per Million (TPM) value for each epigenome and filtered genes to only protein-coding genes located on chromosome 1 through chromosome 22 or chromosome X. We then performed quantile normalization across all epigenomes so that all epigenomes’ TPM values across genes follow the same distribution, reducing the effect of differences between mRNA-seq and total-RNA-seq as well as other experimental differences. After quantile normalization, we performed a log2 transformation after adding a pseudocount of 1 to the gene expression values. In all of the analyses, we used GRCh38 for the genome assembly and GENCODE human release 29 for the gene annotations.

### GECSI method

GECSI takes as input a set of matching chromatin states and gene expression data as a reference compendium, where the samples in the reference compendium are called reference samples, as well as a separate set of gene expression data from new samples. GECSI then outputs chromatin state predictions for the new samples. For each reference sample, GECSI trains a multi-class logistic regression model using features derived from the chromatin state annotations of the *L* nearest reference samples from the reference compendium. The nearest reference samples are determined based on gene expression correlation. After models are trained in the reference compendium, for any given new sample, GECSI generates genome-wide predictions of chromatin states by ensembling the outputs of the models from the *R* nearest reference samples. The predictions are at 200 basepair resolution, where each 200 basepair bin is referred to as a “position.” The steps of GECSI are specified in more detail below.

#### Determining nearest samples

GECSI determines nearest samples to a given sample based on Spearman correlation of gene expression values. Specifically, we denote the transformed gene expression value in sample *M* of gene *g* as *G_M,g_*. Now, we denote the gene expression vector in sample *M* as *G_M_ =* (*G_M,1_,…, G_M,p_*) with a total of *p* genes. The Spearman correlation between two samples, *M* and *N*, is denoted as *C*(*G_M_*, *G_N_*). Here, we denote the set of all samples as *S*. Now, for any given sample *M*, GECSI finds a list of nearest samples *s*_1_,…,*s_k_* that provide the highest values of *C*(*G_M_*, *G_s₁_*),…, *C*(*G_M_*, *G_sₖ_*), where *C*(*G_M_*, *G_s₁_*) > *C*(*G_M_*, *G_s₂_*) >… > *C*(*G_M_*, *G_sₖ_*), among all *C*(*G_M_*, *G_s_*), where *s* ∈ *S* \ *{M}*. If there is a tie, the original ordering based on their position in the input gene expression vector would be preserved.

#### Model training

GECSI trains a multi-class logistic regression model in each reference sample on a random subset of 100,000 positions sampled from the whole genome. For each model, we refer to the sample that the model is trained on as the training sample, denoted as *S*_train_. The chromatin state annotation of *S*_train_ serves as the target that the model learns to predict. The model’s features are obtained from a set of the *L* nearest reference samples to *S*_train_, which are referred to as feature samples. Feature samples are also determined using gene expression information in the same way as described above for determining the nearest samples. GECSI one-hot encodes the chromatin states of the *L* feature samples into binary features, where each feature corresponds to a chromatin state in one sample at a position. In each column, a 1 entry represents the presence of the chromatin state, and 0 means the absence. GECSI uses these binary features to train a multi-class logistic regression model to predict the chromatin state labels of training sample *S*_train_. The model is trained using a lasso penalty with the procedure described below for setting the lasso penalty parameter, *λ*.

#### Model predictions

To make chromatin state predictions for a new sample with gene expression data, *S*_New_, GECSI determines a set of *R* nearest reference samples, *S*_Ref_ = {*S*_Ref_1_,…, *S*_Ref_*R*_ } using the same approach for determining nearest samples as described as above. *R* is a parameter corresponding to the number of models trained for nearest reference samples and included in the model ensemble to generate the final output predictions. GECSI generates chromatin state predictions for a given new sample *S*_New_ by ensembling models trained on *S*_Ref_1_,…, *S*_Ref_*R*_ as defined above. Each of the trained models is used to make a probabilistic prediction for each state and each position. Then, for each state and position, GECSI averages the corresponding prediction probabilities by different models. Finally, for each position GECSI makes a hard assignment of the chromatin state at the position that corresponds to the state that had the highest average probability.

#### Implementation

GECSI’s logistic regression model is implemented using R’s “glmnet” package with the family set as “multinomial”, which fits a multinomial logistic regression model using penalized maximum likelihood. The optimization is performed via the cyclical coordinate descent algorithm. In “glmnet”, the alpha parameter controls the type of regularization: alpha = 1 corresponds to lasso regularization, alpha = 0 to ridge regularization, and values between 0 and 1 correspond to an elastic net combining the two types of regularizations. We set alpha to 1 when calling the glmnet function so that the lasso regularization was used. The lasso regularization parameter was specified by the lambda parameter.

### Cross-validation for evaluation

To evaluate how well predicted chromatin states agree with observed chromatin states, we conducted cross-validation experiments. For GECSI, KNN and Majority State methods, we performed evaluations in 5-fold cross-validation on the 414 fully observed IHEC reference epigenomes. For ChromImpute+ChromHMM, we made predictions in a leave-one-out manner as this is how ChromImpute is designed to make held-out predictions and we expected any effect of this difference would advantage it relative to GECSI.

For each of the folds in the 5-fold cross-validation experiment, approximately 20% of the epigenomes were held out to form a test set used for evaluation. The remaining epigenomes in each fold were further split into a training set and a tuning set, where the tuning set was composed of 6% of the remaining epigenomes. All 414 epigenomes were in exactly one of the test sets. Unless otherwise specified in the following sections, we performed the evaluations on a random sampled set of 200 bp intervals that covered 10% of the whole genome.

We set the values of the hyperparameters of GECSI independently for each fold. For the hyperparameter of the number of trained models to ensemble, *R*, we considered the values: 3, 5, and 10. For the number of features for each individual model, *L*, we considered the values: 5, 10, and 15. For the lasso penalty parameter, *λ*, we considered the values: 0.1, 0.01, 0.001, 0.0001. We performed the selection of parameter values using positions on chromosome 1 in the tuning set of epigenomes. We first tried to select the settings of *λ, R* and *L* based on the combination that optimized median accuracy in each fold. However, we found that such a criteria led to a substantial number of occurrences of specific chromatin states lacking any predicted assignments in specific epigenomes. We observed that the setting of *λ* correlated well with how often this occurred, and thus to reduce the occurrence of this we tuned *λ* based on a different criteria than *R* and *L*. Specifically, we optimized *λ* by selecting the value that maximized the average number of chromatin states that had at least one assigned position in an epigenome in the tuning set across all combinations of values of *R* and *L* that we were considering. After fixing this choice of *λ* we then selected the combination values for *R* and *L* that optimized median accuracy of predictions in the tuning fold **(Supplementary Table 2)**.

### Application on IHEC epigenomes that lack other chromatin state annotations

We applied GECSI to generate chromatin state predictions for all 449 predCS IHEC epigenomes. We used all 414 fullobs epigenomes as the reference data set. For selecting hyperparameter values in this application, we selected values by splitting all fullobs epigenomes into a training and tuning set, with the tuning set composed of 20% of all epigenomes. The selection of values for *R*, *L* and *λ* followed the same procedure as described in the previous section, except not using 5-fold cross-validation, leading to the selection of 5 for *R*, 5 for *L* and 0.0001 for *λ*. Using this set of parameters, we trained GECSI on all fullobs epigenomes in the IHEC compendium and then applied it to generate the chromatin state predictions for the predCS epigenomes.

### Alternative methods and baselines

We compared GECSI to multiple alternative methods and baselines whose implementation is described below.

#### ChromImpute+ChromHMM

We first designed a procedure to impute individual epigenetic tracks from gene expression using ChromImpute^15^ and then from that output generate chromatin state annotations directly with ChromHMM. ChromImpute^15^ takes multiple signal tracks as input and produces imputed signal tracks as output. We note that ChromImpute has been previously used to impute epigenetic and gene expression tracks using other available epigenetic tracks in a sample, but was not previously designed or used to impute epigenetic tracks from gene expression data in a sample. To apply ChromImpute to predict epigenetic tracks based on RNA-seq in a new sample, we first transformed RNA-seq into signal tracks for use with ChromImpute. To do this we annotated all positions within a gene body, defined by the set of positions from the transcription start site to the transcription end site, to the corresponding gene expression value. In cases when multiple genes overlap the same position, we annotated the position with the gene expression value of the gene with the larger coordinate among the smaller one of each gene’s two coordinates. For the expression value we used the log2 transformation of quantile-normalized TPMs, with a pseudocount of 1 added prior to transformation, as described above. We used a resolution of 200 base pairs per bin and all bins that did not intersect the gene body at all were set to 0.

We then applied ChromImpute to impute ChIP-seq data of histone modifications based on this transformed RNA-seq data. The epigenomes that we imputed using ChromImpute were the fullobs epigenomes. The epigenomes that we included in the reference compendium were fullobs epigenomes and epigenomes with observed ChIP-seq data for only a subset of marks considered. For each of the fullobs epigenomes, we imputed six different histone marks: H3K27ac, H3K27me3, H3K36me3, H3K4me1, H3K4me3 and H3K9me3 from RNA-seq using ChromImpute. The imputation of the ChIP-seq signal for one histone mark in a target epigenome was based on its RNA signal track, as well as other reference compendium epigenomes’ RNA signal tracks and ChIP-seq signal tracks of the corresponding histone mark, but not any ChIP-seq data in the target epigenome. We used ChromImpute’s default options except the “-tieglobal” option was used in the “GenerateTrainData” and “Apply” commands and the “-p LOG2RNA” option was used with the “Train” and “Apply” commands where LOG2RNA was the name given to RNA-seq data. ChromImpute models were trained separately for each epigenome and mark combination.

After signal tracks were imputed for the six histone modifications, we then binarized them using the thresholds previously computed for each histone modification for generating ChromHMM annotations on the IHEC epigenomes. These binarization thresholds for imputed data were determined on a mark basis such that the percentage of present calls based on imputed data would be the same on average as based on observed data. The same 18-state ChromHMM model which was used to generate the chromatin state annotations that were used as input to GECSI was used to generate the chromatin state annotations based on the binarization of the imputed histone mark signal.

#### Majority State

Majority state is a baseline method we developed to compare to GECSI. At each position, majority state computed the frequency of each state across the input samples and generated an assignment based on the most frequently appearing state (or the “majority” state) to the corresponding position in a new sample. In cases in which multiple chromatin states share the maximum frequency, we randomly select one of the chromatin states with a maximum frequency.

#### K Nearest Neighbor (KNN)

KNN is an alternative method we developed and to which we also compared GECSI. For each position, the KNN method computed the chromatin state frequency over a set of nearest samples in the same way as GECSI determines its nearest samples as described above. In the case in which multiple chromatin states share the maximum frequency, we randomly selected one of the chromatin states with maximum frequency. We performed parameter selection for the number of nearest samples to use, *k*, across the values 1, 3, 5, 7, 9, for each cross-validation fold, using chromosome 1. The best *k* parameter set was selected according to the accuracy metric **(Supplementary Table 3)**.

#### Closest Reference Sample

We also defined a closest reference sample, which was the sample that had the chromatin state tracks most similar to the target sample in the training data. We computed the most similar reference sample of a target sample by comparing the pairwise agreements between it and all reference samples in the training data. We calculated pairwise agreements as the proportion of the number of positions with matching chromatin state assignments between two samples relative to the total number of positions. The reference sample that had the highest pairwise agreement with the target sample among all reference samples was selected as the closest reference sample. We note that we used the closest reference sample to quantitatively benchmark how well other methods perform in an absolute sense, but since it is defined based on the knowledge of chromatin states in the target sample, it is not an actual imputation method.

### Evaluation metrics

We used multiple evaluation metrics to evaluate GECSI and the alternative methods and baselines.

#### AUPRC

We calculated area under the precision-recall curve (AUPRC), by using precision-recall values that were determined based on ranking predictions using probability assignments from GECSI, state frequencies from the KNN and Majority State baselines, and posterior probabilities from the ChromImpute+ChromHMM method. For each chromatin state, we treated the observed presence or absence of the state at each genomic position as the binary ground truth label. Then we computed the precision-recall curve using a stepwise interpolation, meaning that precision values were assumed to be constant between consecutive recall points. We calculated AUPRC = ∑ *^n^* (*Recall* − *Recall_i_*_−1_) ⋅ *Precision_i_*, where *Recall_i_* = recall at point *i*, *Precision_i_* = precision at point *i*, *n* = the number of all threshold points, with the threshold points defined based on the unique predicted soft assignment values and *Recall_i_* = 0 at *i* = 0.

#### AUROC

As with the AUPRC, we calculated the Area Under the Receiver Operating Characteristic curve (AUROC), using probability assignments from GECSI, state frequencies from KNN and Majority State baselines, and posterior probabilities from the ChromImpute+ ChromHMM method. For each chromatin state, we also treated the observed presence or absence of the state at each genomic position as the binary ground truth label. AUROC was calculated using the pROC package in R, with linear interpolation between unique points defined based on the unique predicted soft assignment values.

#### Accuracy

We calculated the accuracy using the number of positions where the predicted chromatin state label matches the observed chromatin state label, divided by the total number of positions.

#### Weighted Jaccard Average

We calculated the weighted Jaccard average by first calculating Jaccard indices separately for each chromatin state, using the chromatin state hard assignments. For each chromatin state, we calculated the Jaccard index as the ratio of the number of genomic positions overlapping that state in both the predicted and observed tracks (intersection) to the total number of positions assigned to that state in either track (union). We calculated the weighted Jaccard average for each reference epigenome, by first calculating individual Jaccard indices. We calculated Jaccard indices *J_1_*, *J_2_*,…, *J_18_*, corresponding to chromatin states 1, 2,…, 18 respectively. Then, we calculated the weighted Jaccard average *J_avg_* as an average of individual Jaccard indices calculated for each state weighted by the size of the state, or more formally *J_avg_ = ∑^18^_i=1_J_i_s_i_*/*∑^18^_i=1_s_i_*. Here *s_i_* is the size of the *i*^th^ chromatin state, which was defined as the number of genomic positions assigned to that chromatin state averaged over all fully-observed reference epigenomes.

#### Weighted F1 average

We calculated the weighted F1 average by first calculating F1 scores separately for each chromatin state using chromatin state annotation hard assignments. We compared a method’s predictions against the observed chromatin states to first calculate precision and recall, from which an F1 score was then calculated using this formula: F1 = 2 × (*Precision* × *Recall*) / (*Precision* + *Recall*). Then we calculated the weighted F1 average for each method and reference epigenome using individual F1 scores, in the same way as in the weighted Jaccard average as above except that we replaced Jaccard indices with F1 scores.

### Chromatin state genome fraction

For computing a chromatin state genome fraction, for hard assignments, we calculated it as the fraction of the number of genomic positions that were assigned to a state over the total number of genomic positions. For soft assignments, we calculated the fraction as the sum of the predicted probabilities for that state across all positions over the total number of genomic positions.

### Chromatin state grouping

In the analysis of GECSI’s incorrect predictions and the calibration analyses, we used a manual grouping of chromatin state annotations to define similar states. This grouping was based in part on previous biological interpretations of the chromatin states^2^. Specifically, we grouped the chromatin states into the following seven groups:

● Transcription Start Site (TSS) related states: 1_TssA (Active TSS), 2_TssFlnk (Flanking TSS), 3_TssFlnkU (Flanking TSS Upstream), 4_TssFlnkD (Flanking TSS Downstream)
● Transcription related states: 5_Tx (Transcription), 6_TxWk (Weak Transcription)
● Enhancer related states: 7_EnhG1 (Genic enhancer 1), 8_EnhG2 (Genic enhancer 2), 9_EnhA1 (Active Enhancer 1), 10_EnhA1 (Active Enhancer 2), 11_EnhWk (Weak Enhancer)
● States associated with H3K9me3: including 12_ZNF/Rpts (ZNF genes & repeats), 13_Het (Heterochromatin)
● Bivalent states: including 14_TssBiv (Bivalent/Poised TSS) and 15_EnhBiv (Bivalent Enhancer)
● Repressive states associated with H3K27me3: 16_ReprPC (Repressed Polycomb) and 17_ReprPCWk (Weak Repressed Polycomb)
● Quiescent state, which only includes 18_Quies (Quiescent/Low)

### Analysis of GECSI’s incorrect predictions

We characterized GECSI’s incorrect predictions by computing among all genomic positions where the predicted chromatin state annotation did not agree with the observed chromatin state annotation, the proportion of positions where the observed chromatin state was in the same group as the predicted chromatin state. For this analysis we used the chromatin states grouping defined as above and averaged these proportions across the epigenomes. To evaluate whether this proportion was greater than expected by chance, we compared it to an expected proportion, which was the proportion of positions over all non-matching positions where a randomly sampled chromatin state was in the same group as the predicted chromatin state. The chromatin state at a position was randomly sampled according to their relative frequencies in the observed data after excluding the matching chromatin state. In addition, we conducted two similar analyses on incorrect predictions that excluded quiescent regions. In one analysis we removed positions where the predicted chromatin state annotation was quiescent (18_Quies), and for the remaining positions with incorrect predictions we calculated the proportion where the observed chromatin state was in the same group as the GECSI-predicted chromatin state. For this analysis, we calculated the expected proportions by first excluding positions where the predicted annotation was quiescent or where the predicted annotation matched the observed, and then randomly sampling from the chromatin state annotations in remaining positions according to their relative frequencies in observed chromatin states. In the other analysis, we followed the same procedures except that we removed positions where the observed chromatin state annotation was quiescent instead of the predicted.

### Calibration analysis

To evaluate the calibration of predicted probabilities, we used all fullobs IHEC epigenomes and their predicted probabilities for all chromatin states. We first grouped GECSI-predicted probabilities at a randomly sampled 10% of all positions into 50 equally spaced bins spanning the range [0,1]. For each bin, we computed the average predicted probability and the corresponding observed frequency of the target chromatin state. The observed frequency was calculated as the fraction of samples within the bin whose true label matches the predicted state. We then calculated the average Pearson correlation between the observed frequencies and the average predicted probabilities across each combination of epigenome and chromatin state.

We also analyzed the proportions of each observed chromatin state across probability bins of GECSI’s predictions for a specific chromatin state. This was also done across all fullobs IHEC epigenomes and each chromatin state. For this analysis, GECSI-predicted probabilities were divided into 100 equally spaced bins along the range [0,1] and we used all positions in the genome. To reduce instability from small sample sizes and ensure reliable estimation of state composition, we filtered out any bins that had less than 100 positions assigned to it. We specifically evaluated the extent to which GECSI-predicted probabilities of a state being present correlated with a similar state being observed. To do this, for each chromatin state and predicted probability bin, we obtained the fraction of positions with observed state annotations belonging to the same group as the predicted state with the groups defined as previously and positions restricted to those with incorrect predictions. Finally, we computed the Pearson correlation between these fractions and the average predicted probabilities of the bins.

### Gene annotation and CpG island enrichment analysis

We conducted enrichment analyses for GECSI’s predicted chromatin states using external gene annotations and CpG islands. The external gene annotations included exons, gene bodies, transcription start sites (TSSs), transcription end sites (TESs) and regions +/-2kb around TSSs for protein coding genes were from GENCODE v29 annotation^31^. The CpG island annotations were from the UCSC Genome Browser (hg38)^32^. We computed fold enrichment based on all genomic positions using ChromHMM’s OverlapEnrichment command with default parameters. We computed the enrichment for GECSI’s predictions for fullobs epigenomes. We only retained enrichments for a chromatin state in a specific epigenome if its genome coverage was greater than 0.001% to obtain a more robust set of enrichment values. We repeated the analyses also using observed chromatin state annotations. To quantitatively evaluate the consistency of the enrichments between GECSI’s predictions and observed chromatin states, we computed for each annotation and reference epigenome their Pearson correlation restricted to the chromatin state enrichment values that were retained for both. We averaged the correlation values across all reference epigenomes to get an overall correlation score for each gene annotation.

### Transcription factor binding enrichment analysis

We downloaded peak calls of ENCODE TF ChIP-seq experiments^13,33^ from the ENCODE portal^34^. We analyzed data coming from six biosamples: A549, GM12878, HeLa-S3, HepG2, K562, and MCF-7, which were the six cell lines that had data for the greatest number of ENCODE TF ChIP-seq experiments among those represented in the set of fullobs IHEC epigenomes. Specifically, there are 629 experiments from HepG2, 520 experiments from K562, 138 experiments from GM12878, 108 experiments from MCF-7, 58 experiments from A549, and 47 experiments from HeLa-S3. The IHEC IDs for these six cell lines were: IHECRE00001859.4 (A549), IHECRE00001892.4 (GM12878), IHECRE00001851.4 (HeLa-S3), IHECRE00001893.4 (HepG2), IHECRE00001887.4 (K562), and IHECRE00001853.4 (MCF-7).

We computed the enrichments for the GECSI’s predicted chromatin state annotations and observed chromatin state annotations using the same procedures as for the gene annotations and CpG islands above. We also computed separately for each cell line the Pearson correlation between TF binding enrichments using GECSI’s predicted chromatin state annotations and observed chromatin state annotations. Different from how we computed the enrichment correlations above, for this analysis we computed for each cell line correlations based on the concatenation of enrichment values of all TF ChIP-seq experiments concatenated across all chromatin states that were available in both the observed and predicted annotations.

We ordered the TF ChIP-seq experiments based on a previously described ordering method^28^. Specifically, we first performed Pearson correlation among all pairs of the TF ChIP-seq experiments using their enrichment values in all combinations of cell lines and chromatin states. These correlation values were then transformed into a distance function based on the formula *d_i,j_* = 1 - *Cor_i,j_*, where *d_i,j_* represents the distance between TF ChIP-seq experiment *i* and *j*, and *Cor_i,j_* represents the Pearson correlation between the enrichment values of TF ChIP-seq experiment *i* and *j*. Then, we used R’s “seriation” library and “solve_TSP” function to apply a traveling salesman problem solver to minimize the total distance between these TF ChIP-seq experiments’ enrichment values. The solver method is the default heuristic method as provided in the library^35^.

We also calculated TF binding enrichments using GECSI’s predicted chromatin state soft assignments. We computed the fold enrichment using ChromHMM’s OverlapEnrichment command using the “--posterior” option of ChromHMM’s OverlapEnrichment command, with other parameters kept as default. In computing the enrichments, we set all soft assignment values that are below 0.05 to 0 at all positions for all chromatin states. We then renormalized the soft assignments at each position for all chromatin states so that they add up to 1. We used this procedure instead of directly using the soft assignments since we observed that for some states a substantial proportion of the total probability was assigned to locations with small probability and leading to overall weaker enrichments.

### Reference epigenome pairwise correlation and pairwise agreements

To evaluate if chromatin-state patterns across epigenomes in GECSI’s predictions are consistent with those based on observed data, we first calculated pairwise correlations and pairwise agreements between epigenomes for both observed and GECSI-predicted chromatin states using 100,000 randomly sampled positions. For each state, we calculated pairwise Pearson correlation scores for each pair of reference epigenomes. For observed chromatin states, we calculated correlations based on binarized chromatin state assignments. For GECSI-predicted chromatin states, we calculated correlations separately based on binarized chromatin state assignments and soft assignments. For binarized chromatin state assignments, a value of 0 was used to represent an absence of the chromatin state at a position, and 1 the presence. For soft assignments, probability predictions of each chromatin state at each position was used. We also calculated pairwise agreements, which represent an agreement score for each pair of reference epigenomes. An agreement score is the proportion of the number of positions with matching chromatin state annotations between two samples relative to the total number of positions.

To further evaluate the extent to which the agreement and correlation scores reflect biological similarity between reference epigenomes, we split the reference epigenomes into different groups, defined by the harmonized sample intermediate ontology feature in IHEC epigenome metadata v1.2. This led to 47 groups in the fullobs epigenomes, 34 in the predCS epigenomes, and in total 54 when considering both sets of epigenomes. Based on these groups, we then calculated the average pairwise sample correlation for pairs of samples belonging to the same group or different groups separately. We also did the same for pairwise agreement. We calculated these averages using both GECSI-predicted and observed chromatin state annotations in the fullobs epigenomes and for GECSI-predicted annotations in the predCS epigenomes.

For the fullobs epigenomes, we compared pairwise agreements based on GECSI-predicted and those based on observed annotations by how well they can predict whether a pair of reference epigenomes are in the same group. Specifically, we computed the AUPRC using the pairwise agreement scores between reference epigenomes as predictions and epigenome pairs belonging to the same group as the positives and different pairs as the negatives. The precision-recall curve is calculated in the same way as defined in the “Evaluation Metrics” subsection above.

### DNA methylation analysis

For the analysis of average DNA methylation levels in different chromatin states, we used data from WGBS experiments from IHEC, specifically 399 from fullobs epigenomes and 68 from predCS epigenomes. We computed three average values for each chromatin state based on: (1) GECSI’s chromatin state predictions for fullobs epigenomes. (2) GECSI’s chromatin state predictions for predCS epigenomes. (3) Observed chromatin states for fullobs epigenomes. To compute these averages we first computed the average for each epigenome restricted to CpG positions whose DNA methylation value was not considered missing based on having a coverage level (>=3) and then averaged the per-state values across epigenomes.

We also computed the Pearson correlation of average methylation signals across chromatin states. Specifically, we correlated the average GECSI’s predictions in the predCS epigenomes with the average for (1) the observed chromatin state annotations in the fullobs epigenomes and (2) GECSI’s predictions in the fullobs epigenomes.

## Data availability

GECSI-predicted chromatin state tracks along with the input data will be available at the IHEC EpiAtlas portal (https://ihec-epigenomes.org/epiatlas/data/). The TF ChIP-seq enrichments are available from the ENCODE portal with details specified in Supplementary File 2. The gene annotations used for enrichment analysis are available from GENCODE v29 (https://www.gencodegenes.org/human/release_29.html) and CpG islands available from http://hgdownload.soe.ucsc.edu/goldenPath/hg38/database/cpgIslandExt.txt.gz. The code of GECSI can be found at https://github.com/ernstlab/GECSI/ under the open MIT license.

## Supplementary Files

Supplementary File 1: Supplementary Information. Containing Supplementary Figures 1-19 and Supplementary Tables 1-3.

Supplementary File 2: A spreadsheet specifying TF ChIP-seq experiments files. The sheet labeled “TF ChIP-seq downloading links” contains a list of file access URLs associated with all TF ChIP-seq experiments used for enrichment analysis. The sheet labeled “TF ChIP-seq metadata” contains the metadata of TF ChIP-seq experiments used for enrichment analysis. The metadata shown was downloaded from the first URL listed on the sheet “TF ChIP-seq downloading links”.

## Acknowledgements

We thank the International Human Epigenome Consortium (IHEC) for providing access to reprocessed and harmonized epigenomic data comprising the EpiAtlas. We thank Shan Sabri for his assistance on a predecessor project that in part led to this work and Luke Li for helping with track visualization plotting. We thank members of the Ernst Lab for providing helpful discussions and suggestions related to the manuscript.

