## Supplementary File 1 for "GECSI: Large-scale chromatin state imputation from gene expression"

### Contents

Supplementary Figures 1-19
Supplementary Tables 1-3

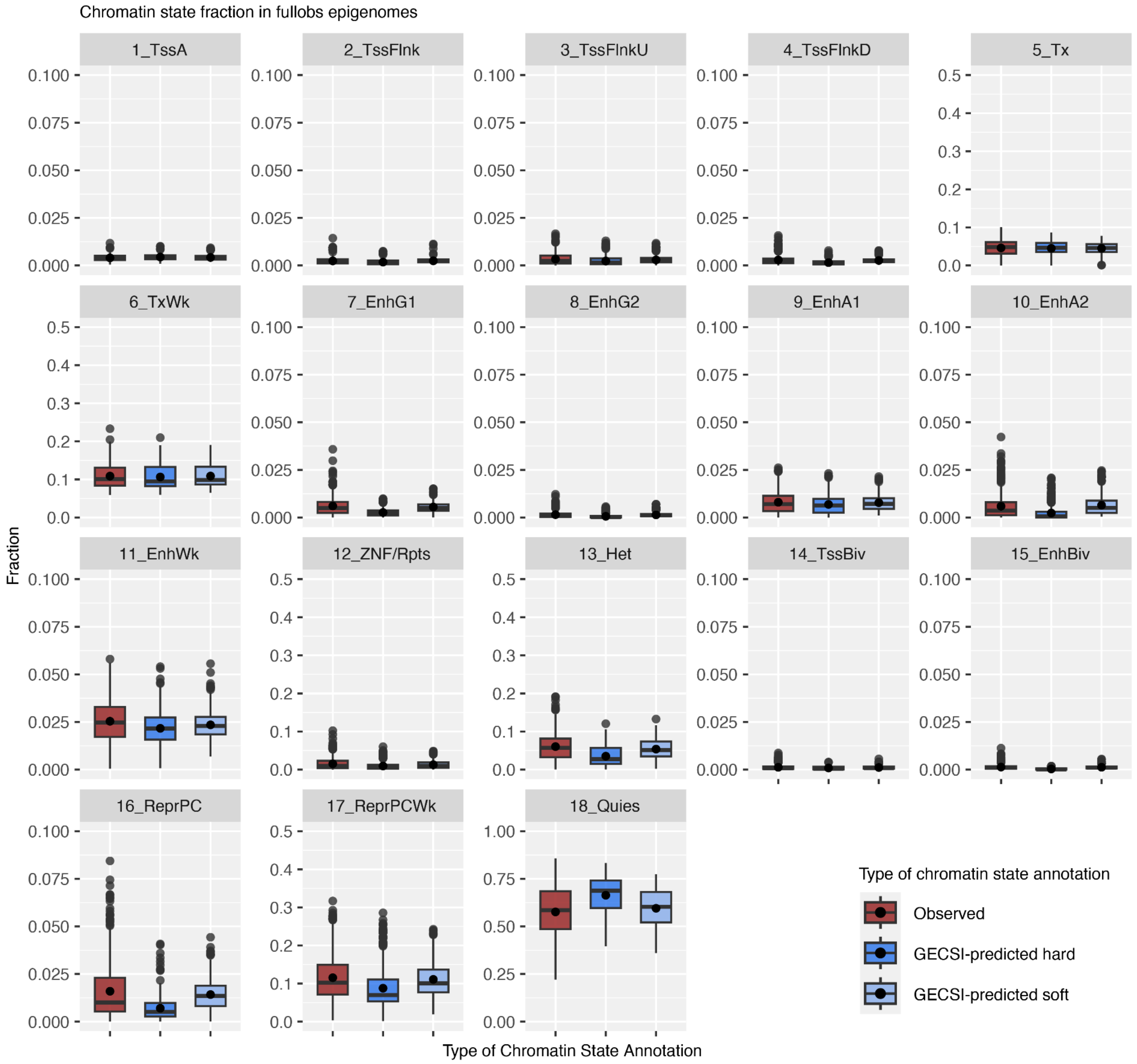


**Supplementary Figure 1: Chromatin state genome coverage fractions for fully observed (fullobs) epigenomes.** Each plot displays for one of the 18 chromatin states the distribution of the genome fraction of the state across all 414 fullobs reference epigenomes with separate boxplots based on observed chromatin state annotations (red), GECSI-predicted chromatin state hard assignments (blue), and GECSI-predicted chromatin state soft assignments (light blue). GECSI predictions were made in cross-validation. For each box, the interquartile range (IQR) is from the 25th to 75th percentile, with the horizontal line inside the box indicating the median, and the dot inside the box indicating the mean. Whiskers extend to the smallest and largest values within 1.5 × IQR from the lower and upper quartiles, respectively; points beyond this range are shown as outliers. Descriptions that correspond to the chromatin state abbreviations are available in Supplementary Table 1.

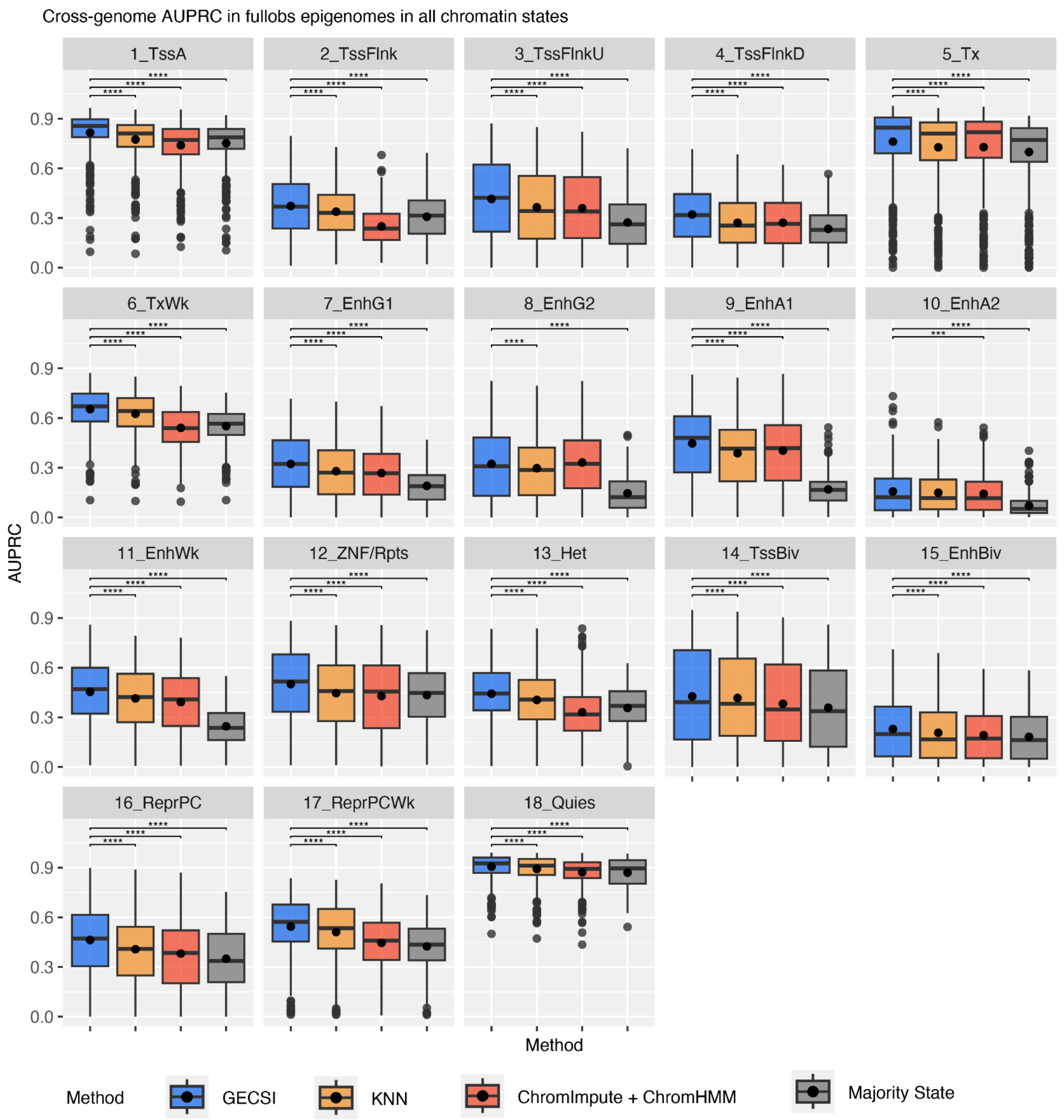


**Supplementary Figure 2: Evaluations on AUPRC scores in the fully observed (fullobs) epigenomes.** Boxplots showing cross-validation results for AUPRC based on soft assignments predicting 18 chromatin states among 414 fullobs reference epigenomes. Each plot displays the AUPRC distribution across all fullobs reference epigenomes for one of the chromatin states. Each box corresponds to one of the methods evaluated: GECSI, KNN, ChromImpute+ChromHMM, or Majority State. Statistical significance between GECSI and other methods was assessed using a one-sided Wilcoxon signed-rank test, and significant differences are indicated with asterisks (*: $p < 0.05$, **: $p < 0.01$, ***: $p < 0.001$, ****: $p < 0.0001$). For each box, the interquartile range (IQR) is from the 25th to 75th percentile, with the horizontal line inside the box indicating the median, and the dot inside the box indicating the mean. Whiskers extend to the smallest and largest values within 1.5 × IQR from the lower and upper quartiles, respectively; points beyond this range are shown as outliers.

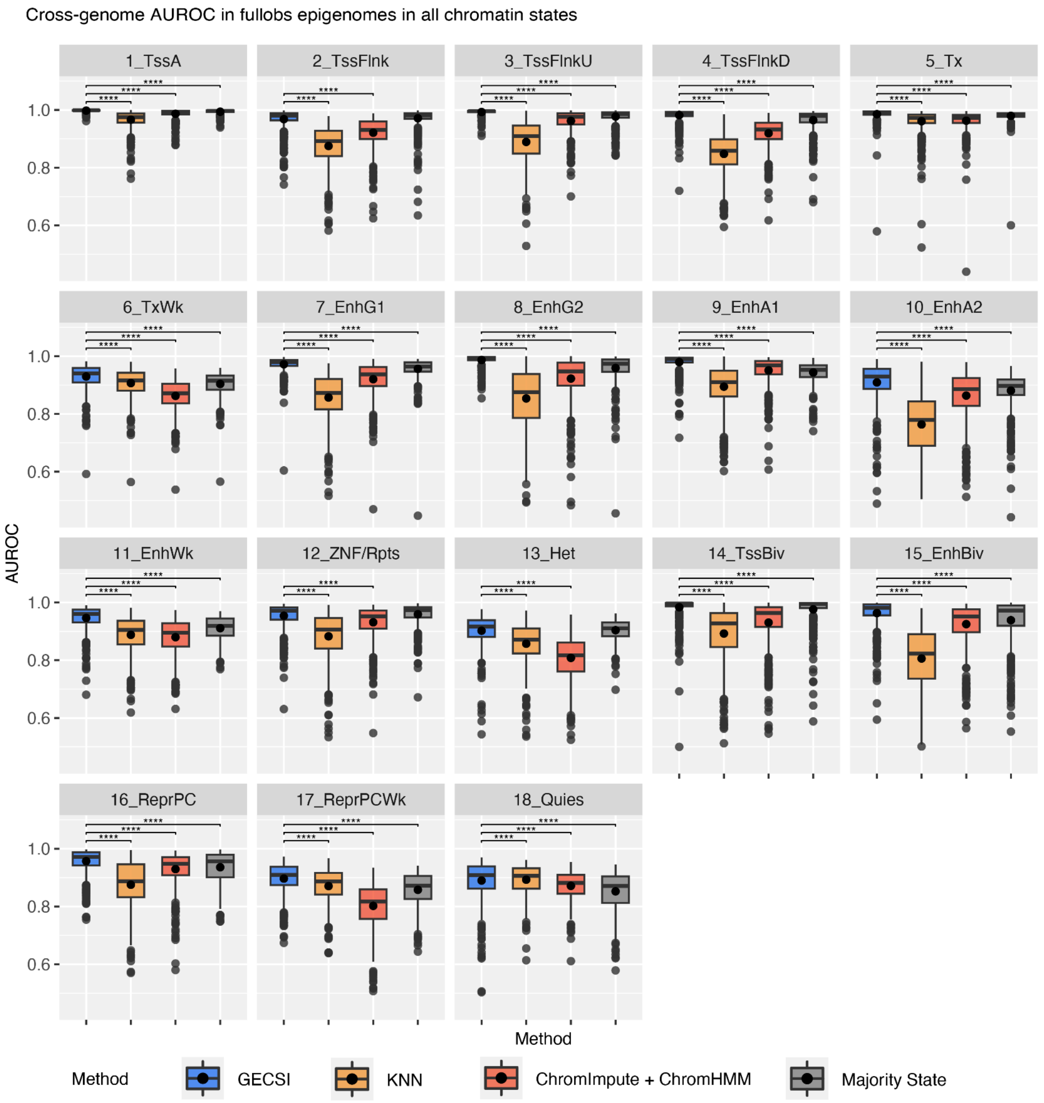


**Supplementary Figure 3: Evaluations on AUROC scores in the fullobs epigenomes.** Boxplots showing cross-validation results for AUROC based on soft assignments predicting 18 chromatin states among 414 fullobs reference epigenomes. Each plot displays the AUROC distribution across all fullobs reference epigenomes for one of the chromatin states. Each box corresponds to one of the methods evaluated: GECSI, KNN, ChromImpute+ChromHMM, and Majority State. Statistical significance between GECSI and other methods was assessed using a one-sided Wilcoxon signed-rank test, and significant differences are indicated with asterisks (*: $p < 0.05$, **: $p < 0.01$, ***: $p < 0.001$, ****: $p < 0.0001$). For each box, the interquartile range (IQR) is from the 25th to 75th percentile, with the horizontal line inside the box indicating the median, and the dot inside the box indicating the mean. Whiskers extend to the smallest and largest values within 1.5 × IQR from the lower and upper quartiles, respectively; points beyond this range are shown as outliers.

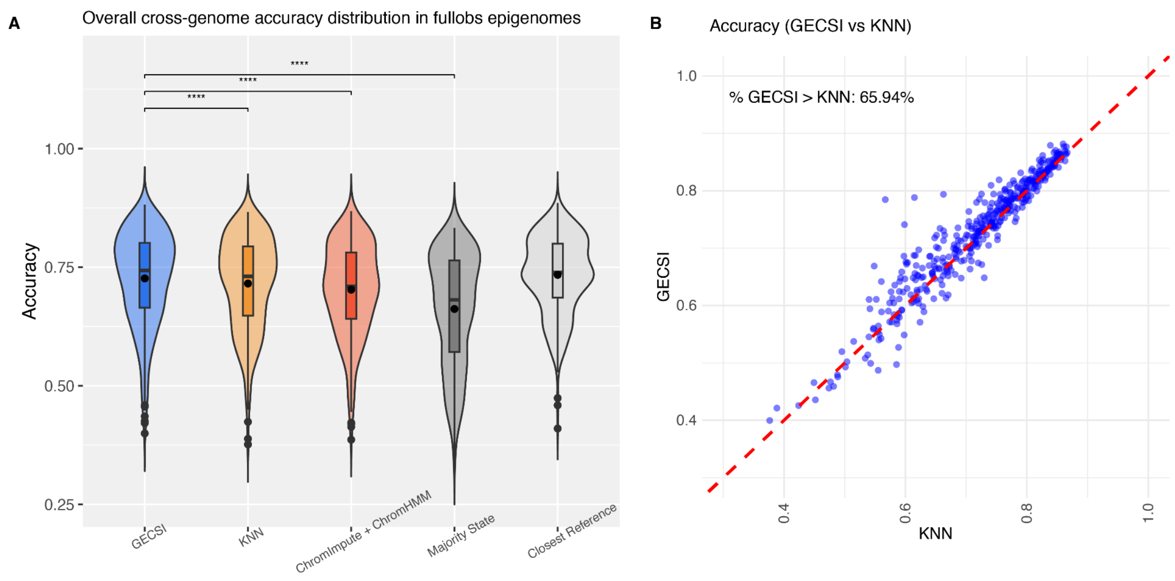


**Supplementary Figure 4: Evaluations on chromatin state prediction accuracy in the fullobs epigenomes.** Shown are cross-validation results for accuracy of predicted hard chromatin state assignments among 414 fullobs reference epigenomes, specifically: **(A)** Violin and boxplots showing the distribution of accuracy across all combinations of epigenomes and chromatin states separately for GECSI, KNN, ChromImpute+ChromHMM, Majority State, and the closest reference benchmark. Statistical significance between GECSI and other methods was assessed using a one-sided Wilcoxon signed-rank test, and significant differences are indicated with asterisks (*: $p < 0.05$, **: $p < 0.01$, ***: $p < 0.001$, ****: $p < 0.0001$). **(B)** Scatterplot comparing accuracy of GECSI method (y-axis) and KNN (x-axis), with the red dashed diagonal line representing y=x. The percentage of points with higher accuracy based on GECSI compared to KNN is shown in the upper left corner of the plot.

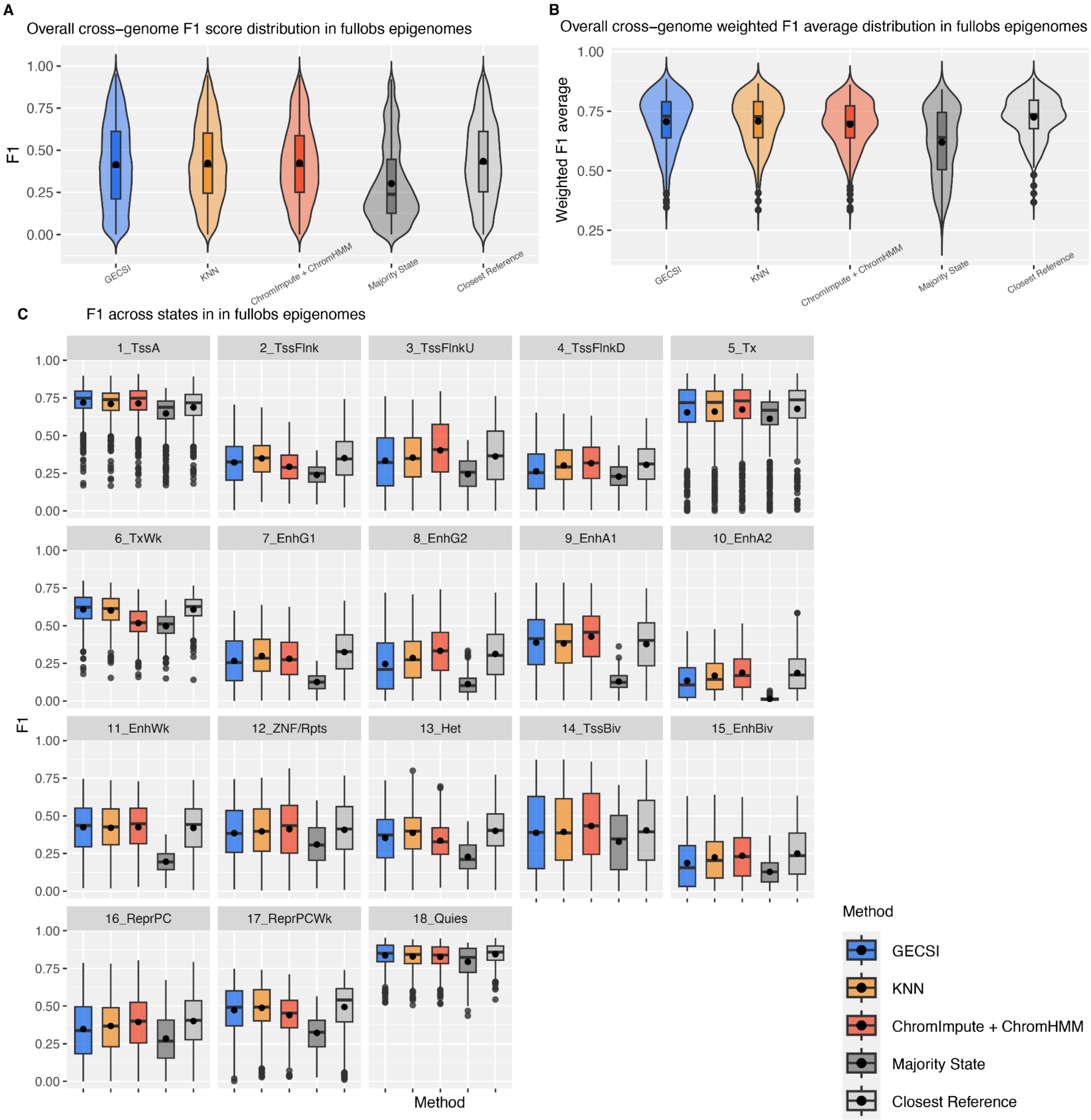


**Supplementary Figure 5: Evaluations on F1 scores and weighted F1 averages in the fullobs epigenomes.** Shown are cross-validation results for F1 score and weighted F1 metrics based on predicted hard chromatin state assignments for 18 chromatin states among 414 fullobs reference epigenomes, comparing GECSI, KNN, ChromImpute+ChromHMM, Majority State, and the closest reference benchmark, specifically: **(A)** Violin and boxplots showing the distribution of F1 scores across all combinations of epigenomes and chromatin states. **(B)** Violin and boxplots showing the distribution of weighted F1 averages across all combinations of epigenomes and chromatin states. **(C)** Boxplots showing the distribution of F1 scores in 18 chromatin states respectively. Each plot displays the corresponding distributions in one chromatin state. For all boxplots, each box represents the interquartile range (IQR) from the 25th to 75th percentile, with the horizontal line inside the box indicating the median, and the dot inside the box indicating the mean. Whiskers extend to the smallest and largest values within 1.5 × IQR from the lower and upper quartiles, respectively; points beyond this range are shown as outliers.

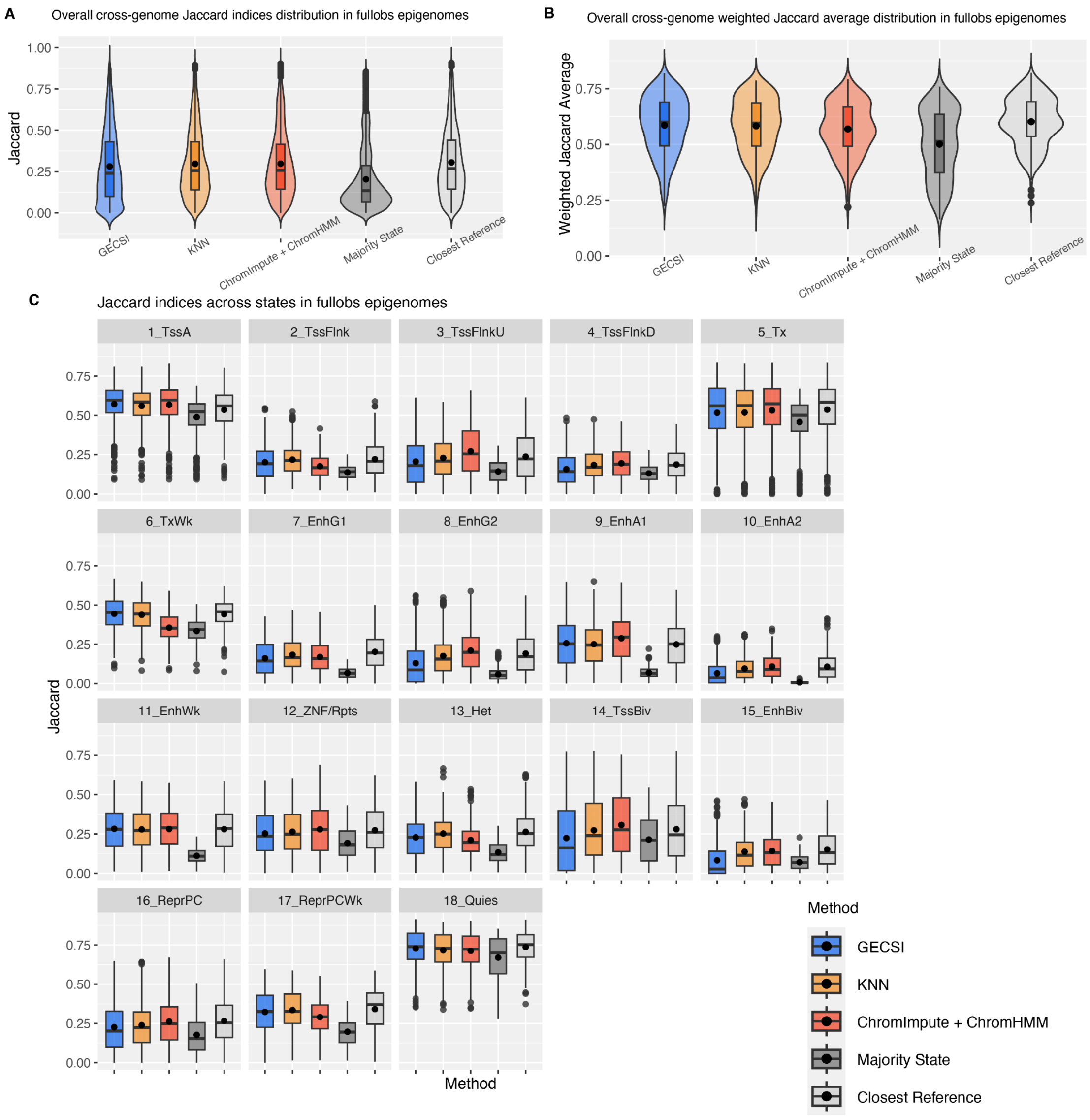


**Supplementary Figure 6: Evaluations on Jaccard indices and weighted Jaccard averages in the fullobs epigenomes.** Shown are cross-validation results for Jaccard index and weighted Jaccard metrics based on predicted hard assignments predictions for 18 chromatin states among 414 fullobs reference epigenomes, comparing GECSI, KNN, ChromImpute+ChromHMM, Majority State, and the closest reference benchmark, specifically: **(A)** Violin and boxplots showing the distribution of Jaccard indices across all combinations of epigenomes and chromatin states. **(B)** Violin and boxplots showing the distribution of weighted Jaccard averages across all combinations of epigenomes and chromatin states. **(C)** Boxplots showing the distribution of Jaccard indices in 18 chromatin states respectively. Each plot displays the corresponding distributions in one chromatin state. For all boxplots, each box represents the interquartile range (IQR) from the 25th to 75th percentile, with the horizontal line inside the box indicating the median, and the dot inside the box indicating the mean. Whiskers extend to the smallest and largest values within 1.5 × IQR from the lower and upper quartiles, respectively; points beyond this range are shown as outliers.

| Predicted \ Observed | 1_TssA | 2_TssFlnk | 3_TssFlnkU | 4_TssFlnkD | 5_Tx | 6_TxWk | 7_EnhG1 | 8_EnhG2 | 9_EnhA1 | 10_EnhA2 | 11_EnhWk | 12_ZNF/Rpts | 13_Het | 14_TssBiv | 15_EnhBiv | 16_ReprPC | 17_ReprPCWk | 18_Quies |
|---|---|---|---|---|---|---|---|---|---|---|---|---|---|---|---|---|---|---|
| 1_TssA | 46101 | 8204 | 7462 | 1330 | 7 | 743 | 23 | 243 | 301 | 1303 | 378 | 32 | 8 | 404 | 31 | 38 | 78 | 273 |
| 2_TssFlnk | 4108 | 12121 | 1536 | 3621 | 9 | 875 | 42 | 80 | 85 | 541 | 952 | 48 | 22 | 1514 | 291 | 197 | 458 | 757 |
| 3_TssFlnkU | 3509 | 1847 | 18275 | 3757 | 8 | 2369 | 121 | 666 | 3950 | 1749 | 2380 | 10 | 11 | 172 | 45 | 19 | 150 | 715 |
| 4_TssFlnkD | 493 | 2623 | 2946 | 10724 | 10 | 1244 | 183 | 256 | 425 | 176 | 3068 | 21 | 14 | 809 | 255 | 63 | 316 | 659 |
| 5_Tx | 16 | 43 | 31 | 88 | 495210 | 132937 | 31482 | 1781 | 354 | 1612 | 1641 | 4858 | 617 | 8 | 19 | 225 | 536 | 8476 |
| 6_TxWk | 649 | 2068 | 1448 | 3334 | 178658 | 999759 | 35577 | 3075 | 13461 | 29795 | 99558 | 8164 | 9988 | 167 | 663 | 2328 | 20006 | 226858 |
| 7_EnhG1 | 16 | 62 | 69 | 414 | 5058 | 3928 | 17916 | 5078 | 864 | 637 | 2510 | 43 | 6 | 3 | 11 | 11 | 22 | 106 |
| 8_EnhG2 | 64 | 62 | 362 | 241 | 183 | 723 | 1788 | 5296 | 612 | 132 | 330 | 2 | 1 | 1 | 1 | 1 | 2 | 23 |
| 9_EnhA1 | 515 | 207 | 8304 | 1537 | 159 | 8305 | 2005 | 4027 | 56194 | 11753 | 21605 | 58 | 99 | 25 | 127 | 65 | 576 | 2833 |
| 10_EnhA2 | 986 | 579 | 1475 | 488 | 355 | 5272 | 1490 | 707 | 5017 | 9601 | 3842 | 41 | 30 | 98 | 101 | 48 | 360 | 1452 |
| 11_EnhWk | 478 | 2017 | 5414 | 16285 | 848 | 48292 | 9116 | 2815 | 34463 | 9420 | 159960 | 329 | 697 | 490 | 2453 | 1048 | 7487 | 30930 |
| 12_ZNF/Rpts | 17 | 50 | 12 | 38 | 2043 | 4551 | 76 | 25 | 73 | 154 | 285 | 95551 | 45664 | 53 | 63 | 2824 | 9065 | 18049 |
| 13_Het | 7 | 38 | 12 | 31 | 355 | 5338 | 19 | 4 | 85 | 163 | 640 | 76026 | 281462 | 16 | 64 | 2741 | 29162 | 128067 |
| 14_TssBiv | 175 | 1040 | 168 | 987 | 2 | 42 | 2 | 2 | 5 | 45 | 263 | 25 | 6 | 6736 | 1368 | 1260 | 369 | 148 |
| 15_EnhBiv | 17 | 64 | 38 | 159 | 5 | 39 | 11 | 3 | 35 | 14 | 374 | 25 | 10 | 981 | 2351 | 790 | 256 | 48 |
| 16_ReprPC | 64 | 394 | 53 | 186 | 162 | 1330 | 64 | 15 | 79 | 78 | 990 | 2015 | 1531 | 1876 | 4436 | 58421 | 33903 | 5751 |
| 17_ReprPCWk | 141 | 1041 | 203 | 746 | 972 | 22475 | 256 | 52 | 921 | 1723 | 10993 | 12378 | 45521 | 1174 | 5636 | 127327 | 804309 | 346950 |
| 18_Quies | 931 | 5461 | 2092 | 5455 | 17126 | 489398 | 2749 | 385 | 12040 | 20367 | 116590 | 67516 | 577053 | 1077 | 4176 | 54005 | 930822 | 7591249 |

**Supplementary Figure 7: Confusion matrix between observed and GECSI-predicted chromatin states in the fullobs epigenomes.** A confusion matrix showing the average number of positions which are assigned to each combination of GECSI-predicted and observed chromatin states across the whole genome. Each value shows the average number of positions across all positions in the fullobs epigenomes with the corresponding GECSI-predicted (rows) and observed (columns) chromatin states. The color scale is column specific and reflects the individual values normalized by the sum of the values in the column. Darker green reflects a higher proportion while white reflects low proportion. Chromatin states are ordered and colored as previously[1]. Descriptions that correspond to the chromatin state abbreviations are available in Supplementary Table 1.

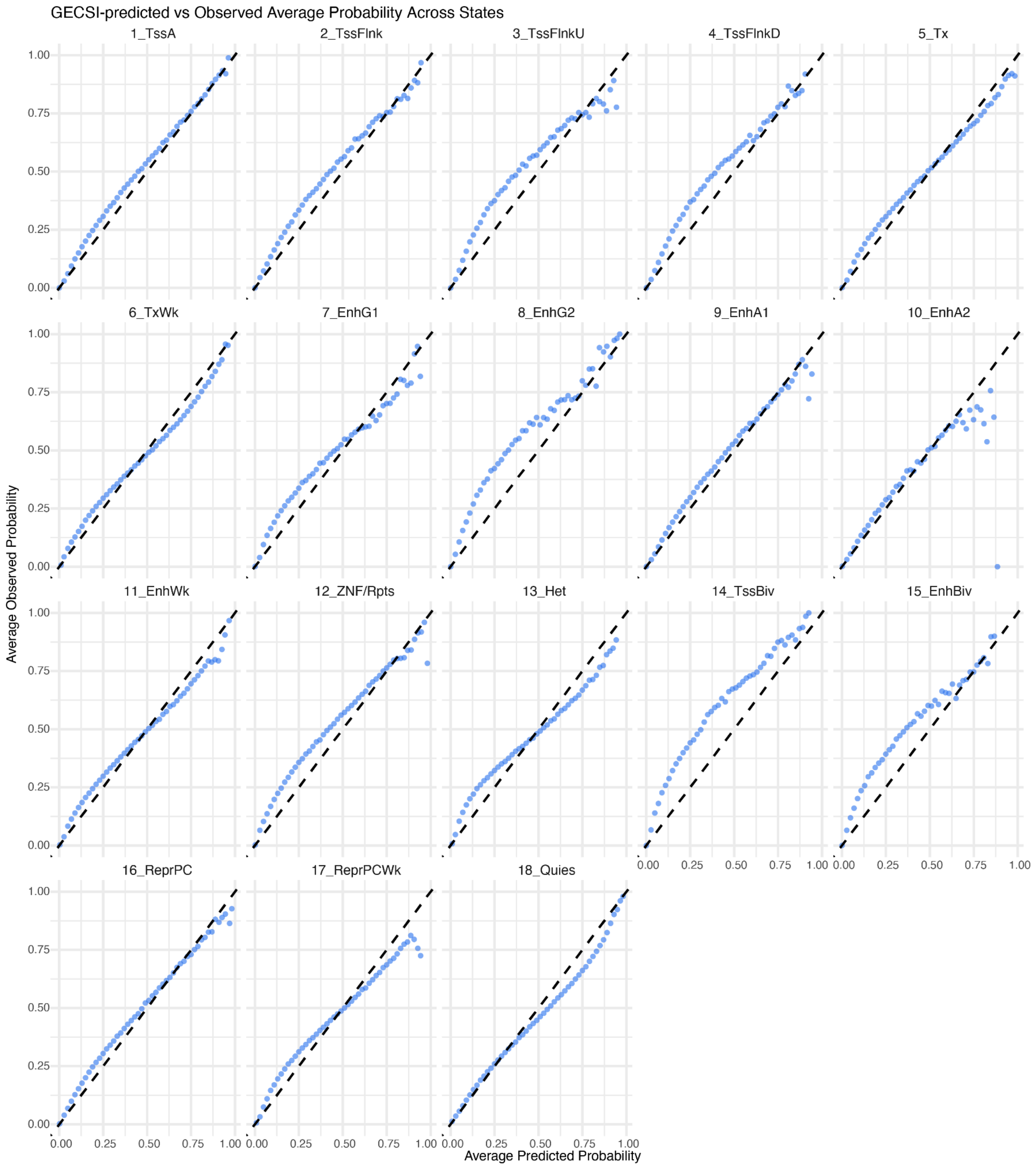


**Supplementary Figure 8: Calibration analysis of GECSI-predicted probabilities in 18 chromatin states**. Scatterplots comparing average GECSI-predicted probabilities for a chromatin state (x-axis) to the average observed frequency of the state (y-axis) from a cross-validation evaluation among 414 fullobs reference epigenomes. Each plot displays the values for one out of the 18 chromatin states. For each chromatin state and epigenome, genomic positions were grouped based on the predicted probability of the state in the epigenome and divided into 50 equal sub-intervals of the interval [0,1]. Average predicted probabilities and average observed probabilities are computed across all positions in each group and all epigenomes. The diagonal black dashed line marks the y=x line, representing perfect calibration.

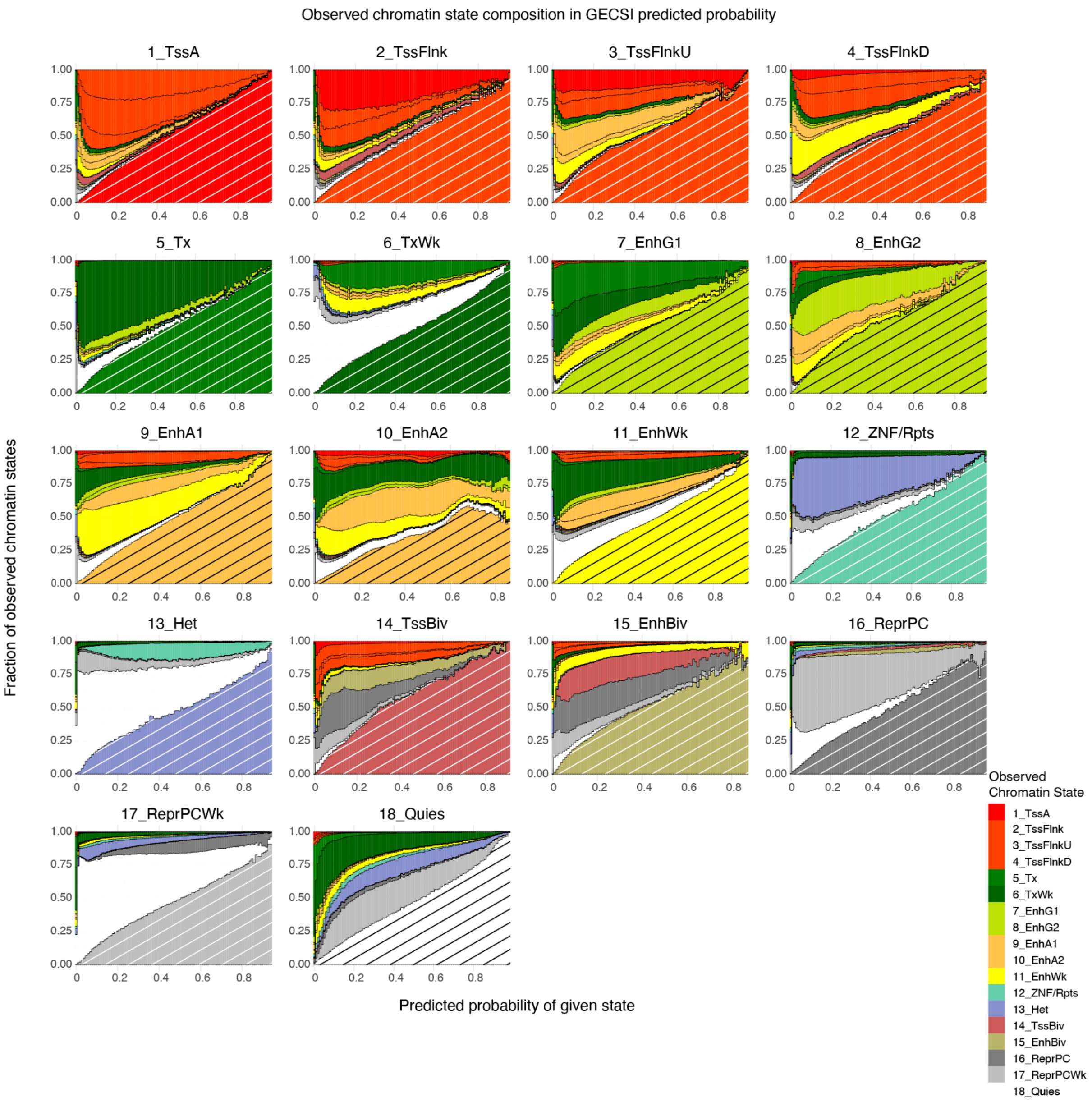


**Supplementary Figure 9: Visualization of observed chromatin state composition with respect to GECSI-predicted probabilities for the 18 chromatin states in the fullobs epigenomes**. Each plot shows the distribution of the observed chromatin states conditioned on GECSI's predicted probabilities of a chromatin state, which are grouped into 100 bins based on the predicted probability of the given state shown on the x-axis. Only bins that had more than or equal to 100 positions assigned to it are displayed and the rest are filtered out and omitted as they only occur at the ends of the available probability ranges. The bars that represent the fractions of a chromatin state are colored by the color-coding of that state. For each plot, the color bar corresponding to the given state is located at the bottom of each bar with diagonal shading overlaid for better visualization. The remaining chromatin states are stacked above it, following the standard order of the chromatin states.

Observed

| State | CpG sites | Exons | Genes | TES | TSS | TSS 2kb |
|---|---|---|---|---|---|---|
| 1_TssA | 73.8 | 10.3 | 1.8 | 1.9 | 71.0 | 12.5 |
| 2_TssFlnk | 49.1 | 6.4 | 1.5 | 1.9 | 40.2 | 10.4 |
| 3_TssFlnkU | 15.8 | 4.5 | 1.7 | 2.8 | 10.8 | 8.7 |
| 4_TssFlnkD | 15.6 | 4.5 | 1.7 | 3.0 | 9.7 | 9.3 |
| 5_Tx | 0.8 | 5.2 | 2.3 | 6.8 | 2.4 | 3.7 |
| 6_TxWk | 0.3 | 1.8 | 1.9 | 1.9 | 0.9 | 1.8 |
| 7_EnhG1 | 1.0 | 5.6 | 2.2 | 8.5 | 3.1 | 4.1 |
| 8_EnhG2 | 1.8 | 6.9 | 2.2 | 11.4 | 4.4 | 5.3 |
| 9_EnhA1 | 0.2 | 1.3 | 1.5 | 1.3 | 1.4 | 2.1 |
| 10_EnhA2 | 0.9 | 1.3 | 1.4 | 1.3 | 1.7 | 2.2 |
| 11_EnhWk | 0.5 | 1.2 | 1.5 | 1.1 | 1.2 | 2.2 |
| 12_ZNF/Rpts | 0.8 | 1.6 | 0.9 | 1.2 | 0.4 | 0.6 |
| 13_Het | 0.2 | 0.2 | 0.6 | 0.2 | 0.1 | 0.2 |
| 14_TssBiv | 88.1 | 8.4 | 1.5 | 2.1 | 26.0 | 10.1 |
| 15_EnhBiv | 34.3 | 4.4 | 1.3 | 2.5 | 8.2 | 6.3 |
| 16_ReprPC | 4.7 | 1.7 | 1.0 | 1.3 | 1.8 | 2.8 |
| 17_ReprPCWk | 0.4 | 0.7 | 0.9 | 0.6 | 0.5 | 0.8 |
| 18_Quies | 0.2 | 0.4 | 0.8 | 0.4 | 0.2 | 0.4 |

GECSI predicted

| State | CpG sites | Exons | Genes | TES | TSS | TSS 2kb |
|---|---|---|---|---|---|---|
| 1_TssA | 74.7 | 10.3 | 1.8 | 1.9 | 70.7 | 12.7 |
| 2_TssFlnk | 67.8 | 7.3 | 1.6 | 1.9 | 46.8 | 11.4 |
| 3_TssFlnkU | 11.6 | 4.6 | 1.7 | 3.0 | 9.5 | 9.5 |
| 4_TssFlnkD | 18.2 | 5.1 | 1.8 | 3.2 | 10.7 | 11.0 |
| 5_Tx | 0.8 | 5.5 | 2.3 | 7.4 | 2.6 | 3.9 |
| 6_TxWk | 0.4 | 1.9 | 2.0 | 2.0 | 0.9 | 1.9 |
| 7_EnhG1 | 1.1 | 6.6 | 2.3 | 10.9 | 3.8 | 4.8 |
| 8_EnhG2 | 1.9 | 8.0 | 2.2 | 13.7 | 5.1 | 6.0 |
| 9_EnhA1 | 0.2 | 1.4 | 1.5 | 1.4 | 1.4 | 2.2 |
| 10_EnhA2 | 2.5 | 1.7 | 1.4 | 1.8 | 3.1 | 3.1 |
| 11_EnhWk | 0.6 | 1.3 | 1.5 | 1.3 | 1.4 | 2.6 |
| 12_ZNF/Rpts | 0.9 | 2.4 | 1.1 | 1.9 | 0.6 | 0.8 |
| 13_Het | 0.3 | 0.3 | 0.5 | 0.2 | 0.1 | 0.2 |
| 14_TssBiv | 101.5 | 9.2 | 1.6 | 2.2 | 29.7 | 10.8 |
| 15_EnhBiv | 64.4 | 6.2 | 1.4 | 2.9 | 12.9 | 8.2 |
| 16_ReprPC | 10.9 | 2.5 | 1.1 | 1.6 | 3.1 | 4.5 |
| 17_ReprPCWk | 0.6 | 0.8 | 0.9 | 0.7 | 0.6 | 1.0 |
| 18_Quies | 0.2 | 0.4 | 0.8 | 0.4 | 0.3 | 0.4 |

**Supplementary Figure 10: External genomic annotation and CpG island enrichments in different GECSI-predicted chromatin states across the fullobs epigenomes.** Heatmaps showing chromatin-state-specific enrichments of different external categories based on observed chromatin state annotations (left) and GECSI's predictions (right). Each column corresponds to an external category, specifically CpG islands, exons, gene bodies (Genes), transcription end sites (TES), transcription start sites (TSS) and 2kb around TSS (TSS 2kb). Each row represents a chromatin state as shown in the first column of color labels. Each grid represents a median enrichment value of the corresponding chromatin state and external category across all reference epigenomes. The coloring scheme is based on enrichment values normalized between 0 and 1 for each column separately, where the maximum is scaled to 1 and the minimum to 0. Specifically, yellow represents higher normalized enrichments and navy blue represents lower normalized enrichments.

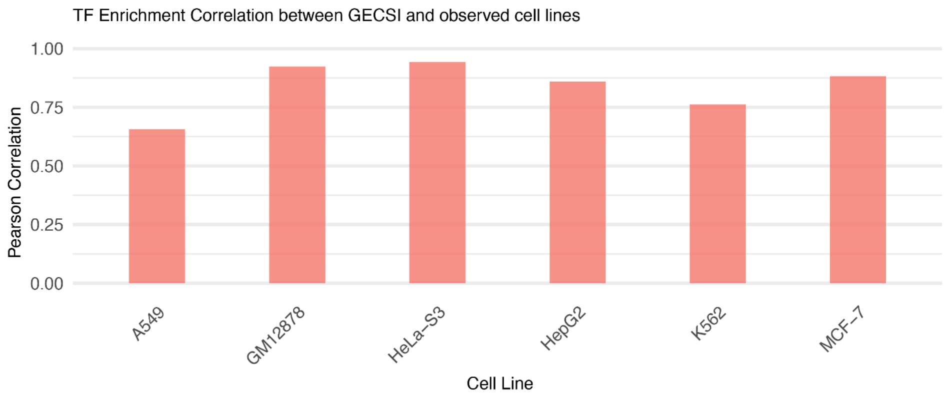


**Supplementary Figure 11: TF binding enrichment correlation bar plot.** Barplot showing for each cell line the correlation between GECSI's predicted chromatin states and observed chromatin states, based on their fold enrichments computed using peak calls from TF ChIP-seq experiments. Each bar represents a Pearson correlation score for a cell line, where the correlation is calculated between the fold enrichments computed in observed chromatin states and the fold enrichments computed in GECSI-predicted chromatin states.

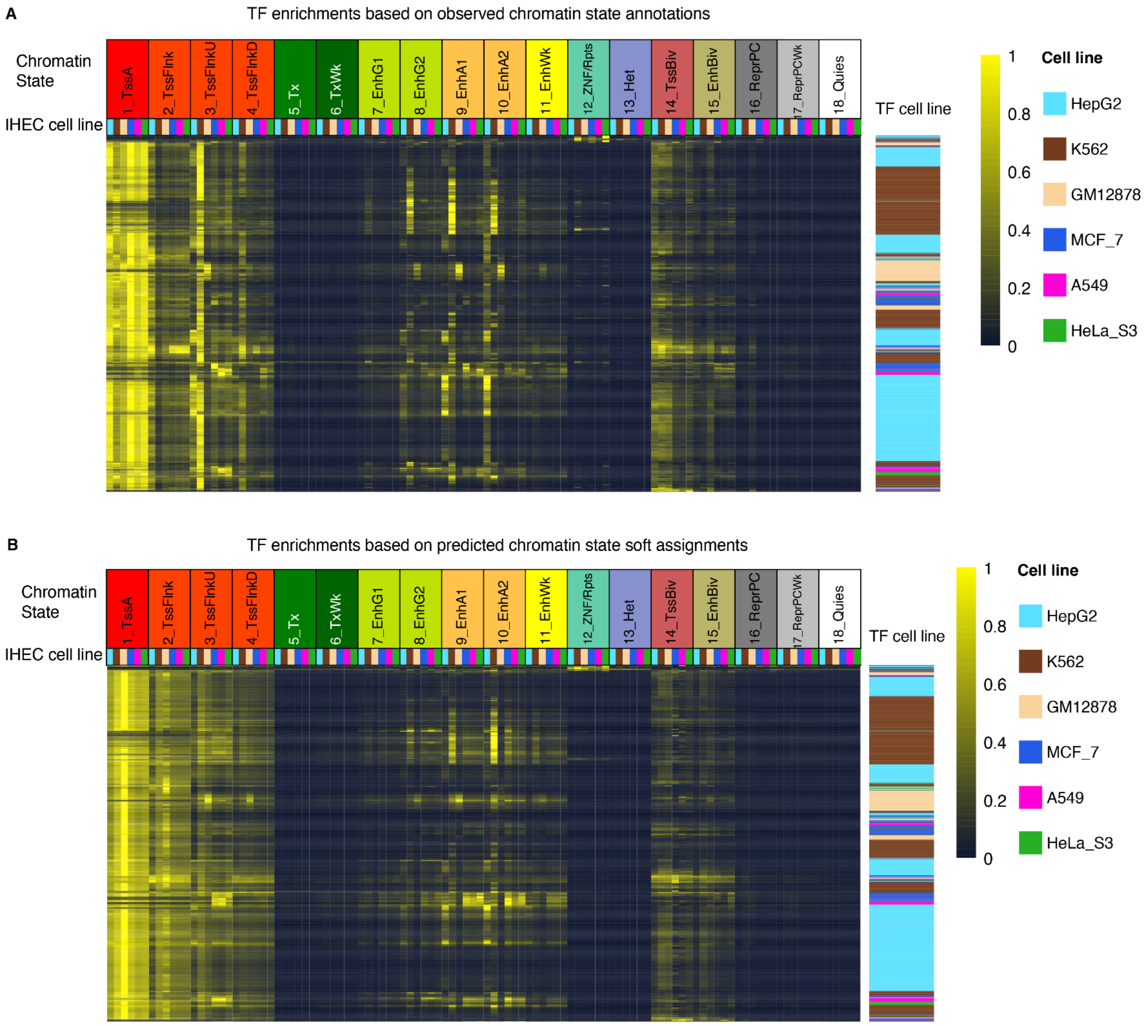


**Supplementary Figure 12: Transcription factor (TF) binding enrichments in observed chromatin annotations and GECSI-predicted chromatin state soft assignments across six cell lines. (A)** TF binding enrichment heatmap with fold enrichments calculated using peak calls from TF ChIP-seq experiments and observed chromatin state hard assignments. **(B)** TF binding enrichment heatmap with the fold enrichments calculated using peak calls from TF ChIP-seq experiments and GECSI-predicted chromatin state soft assignments with low probabilities (<0.05) truncated to zero and renormalized (Methods). For both (A) and (B), each row corresponds to a TF ChIP-seq experiment in a given cell line. Each column corresponds to a chromatin state-cell line combination. The entries in the heatmap are fold enrichments normalized between 0 and 1 for each row separately, where the maximum is scaled to 1 and the minimum to 0. The top right color scale shows the scaled enrichments with yellow representing higher enrichments and blue representing lower enrichments. The row ordering of TF ChIP-seq experiments for (A) and (B) were obtained by applying a traveling salesman problem solver to the pairwise correlations of the chromatin state enrichments values in (A) (Methods)[2] and the same ordering was then applied in (B). The columns are ordered first by chromatin states and then by individual cell lines using the following order: HepG2, K562, GM12878, MCF-7, A549, and HeLa-S3. The top row of labels corresponds to chromatin states and the second row of color labels corresponds to cell lines. The rows are color-coded by the cell line of the TF ChIP-seq experiment, with the same color-coding as the column labels.

| State | Pearson |
|---|---|
| 1_TssA | 0.443 |
| 2_TssFlnk | 0.519 |
| 3_TssFlnkU | 0.581 |
| 4_TssFlnkD | 0.498 |
| 5_Tx | 0.546 |
| 6_TxWk | 0.569 |
| 7_EnhG1 | 0.665 |
| 8_EnhG2 | 0.662 |
| 9_EnhA1 | 0.773 |
| 10_EnhA2 | 0.728 |
| 11_EnhWk | 0.771 |
| 12_ZNF/Rpts | 0.496 |
| 13_Het | 0.507 |
| 14_TssBiv | 0.580 |
| 15_EnhBiv | 0.487 |
| 16_ReprPC | 0.621 |
| 17_ReprPCWk | 0.658 |
| 18_Quies | 0.745 |

**Supplementary Figure 13: Pearson correlation between pairwise sample correlation values derived from observed and GECSI-predicted chromatin states in the fullobs epigenomes.** The figure shows for each chromatin state the Pearson correlation between pairwise sample correlations computed based on observed and GECSI's predicted chromatin states. Pairwise sample correlation scores were calculated in the fullobs epigenomes based on the binary presence or absence of the state.

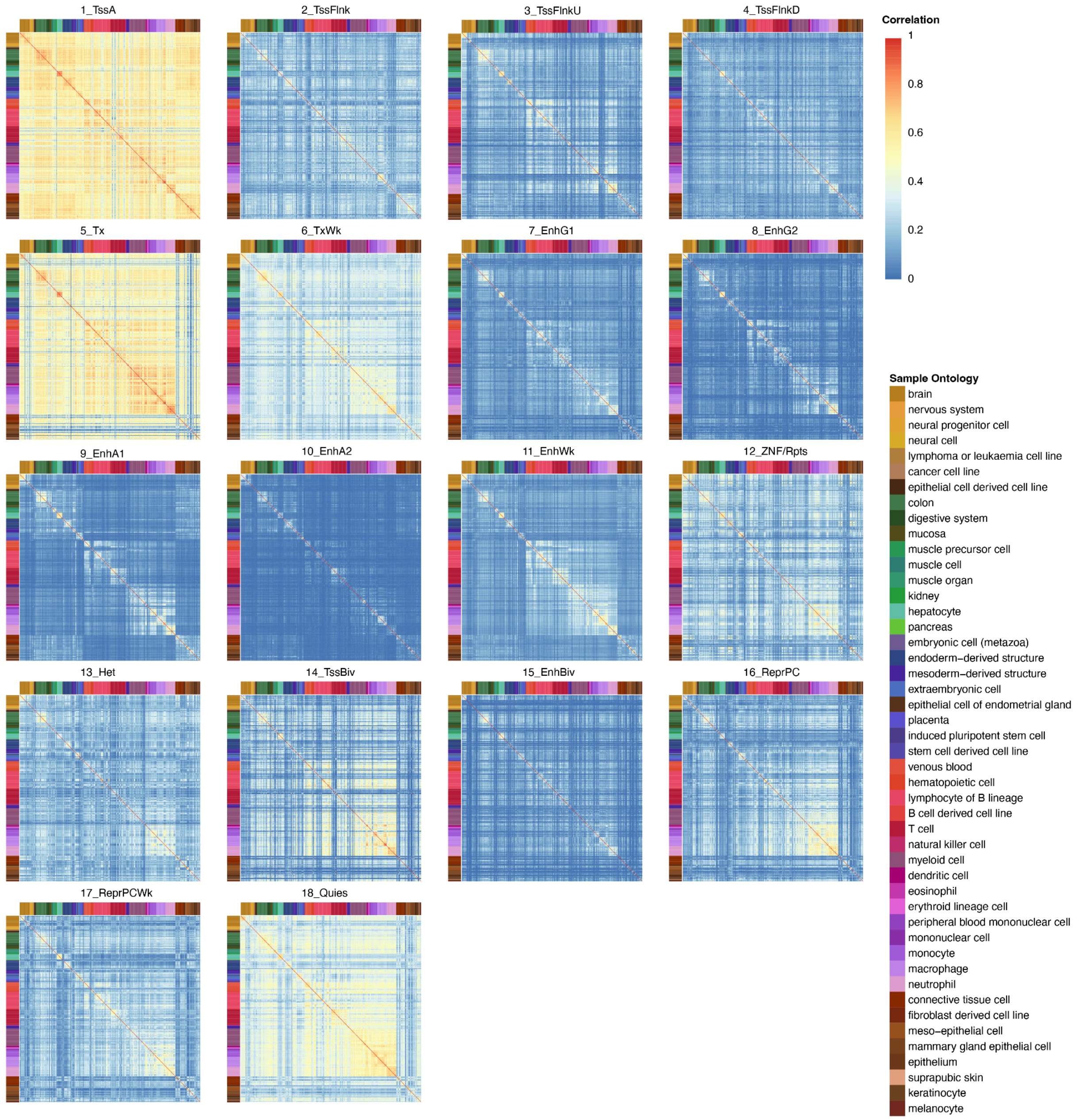


**Supplementary Figure 14: Pairwise sample correlation generated from observed chromatin state annotations in 18 chromatin states in the fullobs samples.** Heatmaps showing for each of the 18 chromatin states the pairwise correlation calculated using the presence or absence of the observed chromatin state among 414 fullobs IHEC epigenomes. In each heatmap, each row and column represents an epigenome, and each entry represents a Pearson correlation value between the epigenome of the row and the epigenome of the column. Only epigenomes with a corresponding chromatin state assignment are shown. The color of the entries follows the color scale on the top right of the figure. Correlation results are ordered based on the provided sample ordering of IHEC, and color annotation of each sample is based on the sample intermediate ontology in the IHEC metadata using the IHEC coloring.

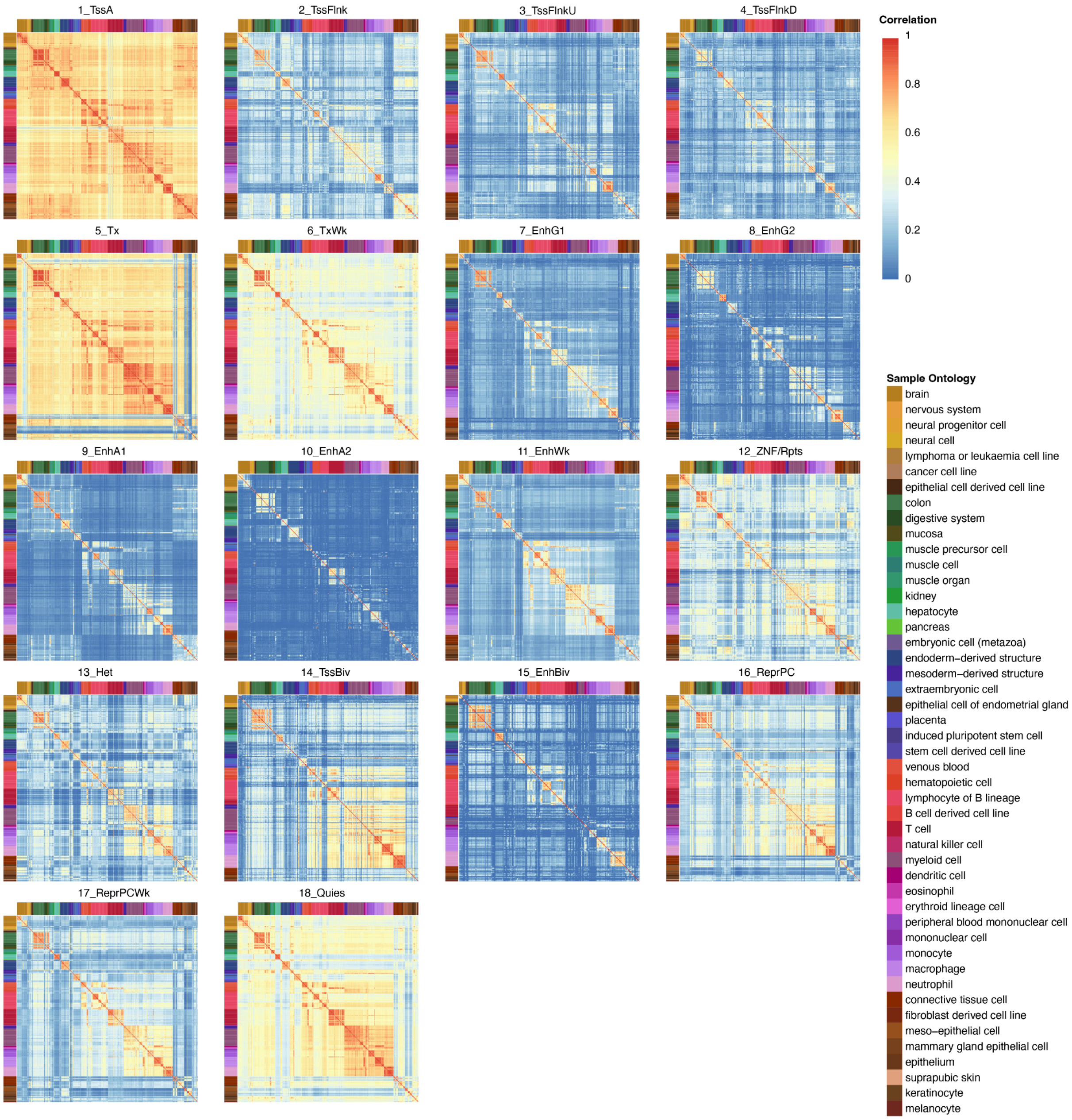


**Supplementary Figure 15: Pairwise sample correlation generated from GECSI-predicted chromatin state annotations in 18 chromatin states in the fullobs samples.** Heatmaps showing for each of the 18 chromatin states the pairwise correlation calculated using the presence or absence of the GECSI-predicted chromatin state among 414 fullobs IHEC epigenomes. In each heatmap, each row and column represents an epigenome, and each entry represents a Pearson correlation value between the epigenome of the row and the epigenome of the column. Only epigenomes with a corresponding chromatin state assignment are shown. The color of the entries follows the color scale on the top right of the figure. Correlation results are ordered based on the provided sample ordering of IHEC, and color annotation of each sample is based on the sample intermediate ontology in the IHEC metadata using the IHEC coloring.

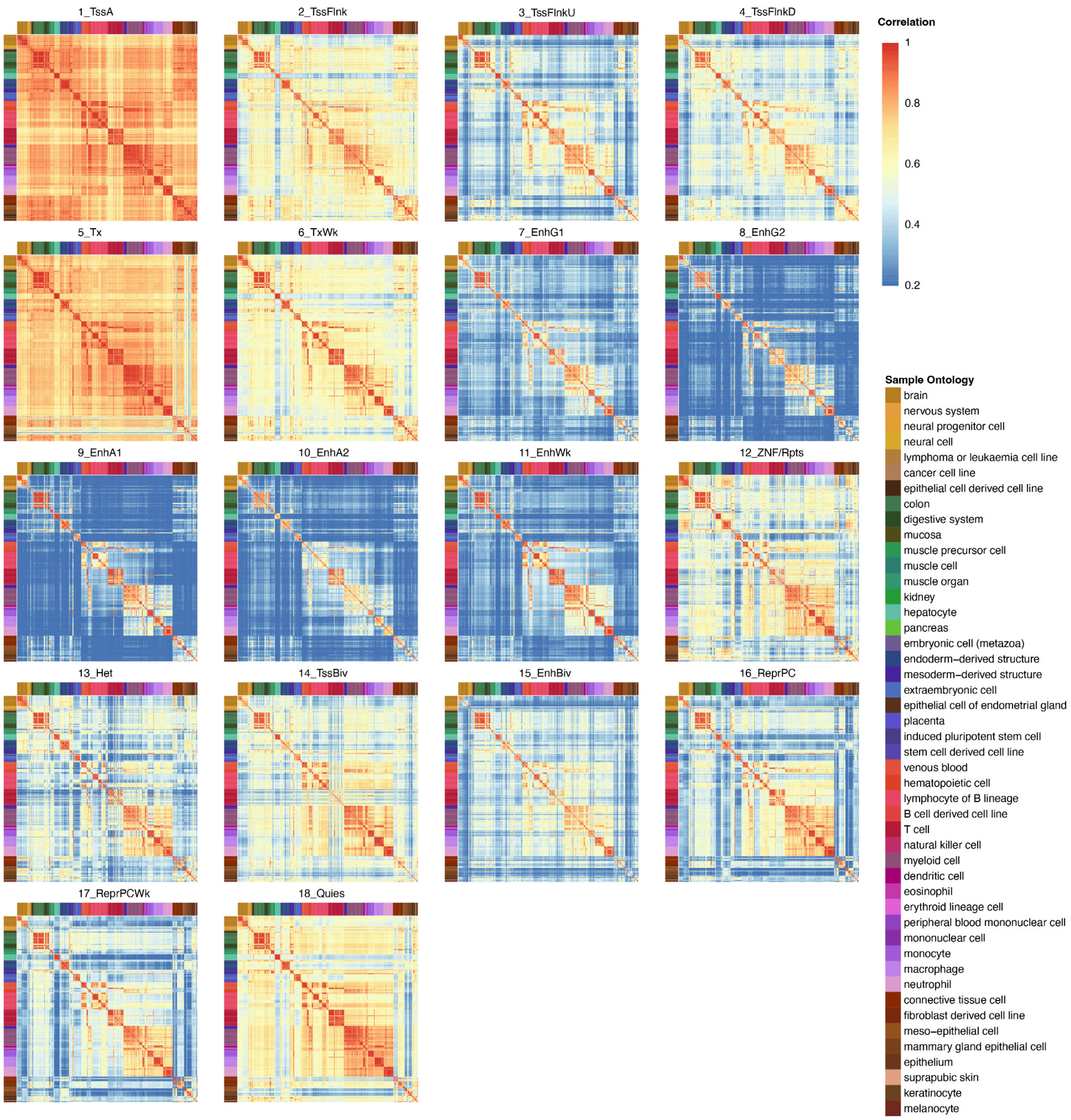


**Supplementary Figure 16: Pairwise sample correlation generated from GECSI-predicted chromatin state soft assignments in 18 chromatin states in the fullobs epigenomes.** Heatmaps showing for each of the 18 chromatin states the pairwise correlation calculated using the soft assignments of the GECSI-predicted chromatin state among the 414 fullobs IHEC epigenomes. In each heatmap, each row and column represents an epigenome, and each entry represents a Pearson correlation value between the epigenome of the row and the epigenome of the column. The color of the entries follows the color scale on the top right of the figure (0.2–1.0), with values below 0.2 mapped to the color for 0.2. Correlation results are ordered based on the provided sample ordering of IHEC, and color annotation of each sample is based on the sample intermediate ontology in the IHEC metadata using the IHEC coloring.

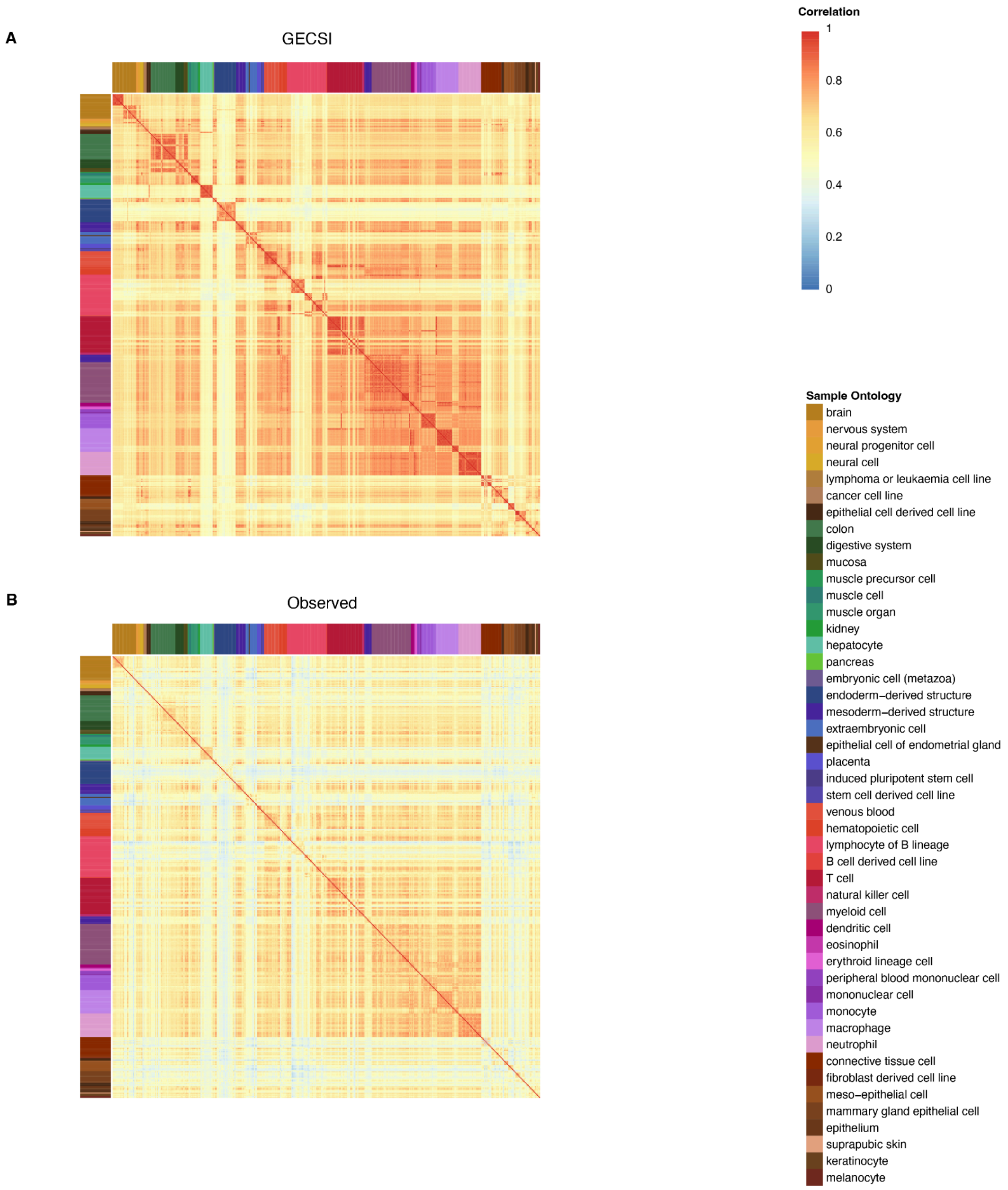


**Supplementary Figure 17: Pairwise sample agreements generated from GECSI, KNN, ChromImpute+ChromHMM and observed chromatin states.** Heatmaps showing the pairwise agreements calculated using **(A)** GECSI-predicted and **(B)** observed chromatin states among the 414 fullobs IHEC epigenomes. Each row and column represents an epigenome, and each value represents an agreement score between the epigenome of the row and the epigenome of the column. The color of the entries follows the color scale on the top right of the figure. Correlation results are ordered based on provided sample ordering of IHEC, and color annotation of each sample is based on sample intermediate ontology in the IHEC metadata using the color codes provided by IHEC.

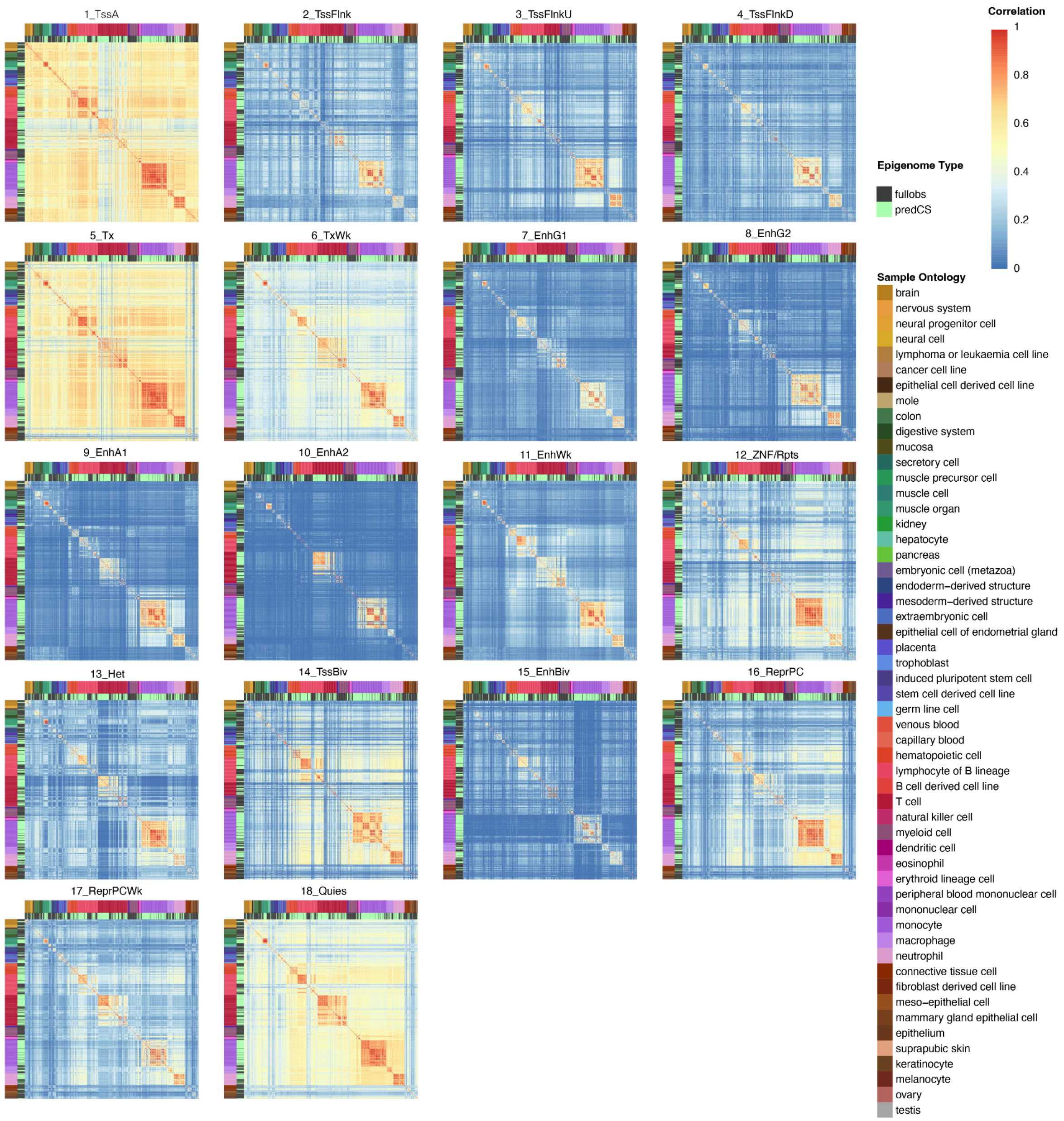


**Supplementary Figure 18: Pairwise sample correlation generated from observed + GECSI imputed chromatin states in 18 chromatin states in the fullobs and predCS epigenomes.** Heatmaps showing for each of the 18 chromatin states the pairwise correlation calculated using the binary presence or absence of the observed chromatin states among the 414 fullobs IHEC epigenomes and GECSI-predicted chromatin states among the 449 predCS epigenomes. In each heatmap, each row and column represents an epigenome, and each entry represents a Pearson correlation between the epigenome of the row and the epigenome of the column. Only epigenomes with a corresponding chromatin state assignment are shown. The color of the entries follows the color scale shown on the top right. Correlation results are ordered based on sample ordering provided by the IHEC. The outer multiple-color annotation of epigenomes is based on sample intermediate ontology in the IHEC metadata using the IHEC coloring, while the inner binary color annotation is based on whether the chromatin state is observed (in the fullobs epigenomes) or GECSI-predicted (in the predCS epigenomes).

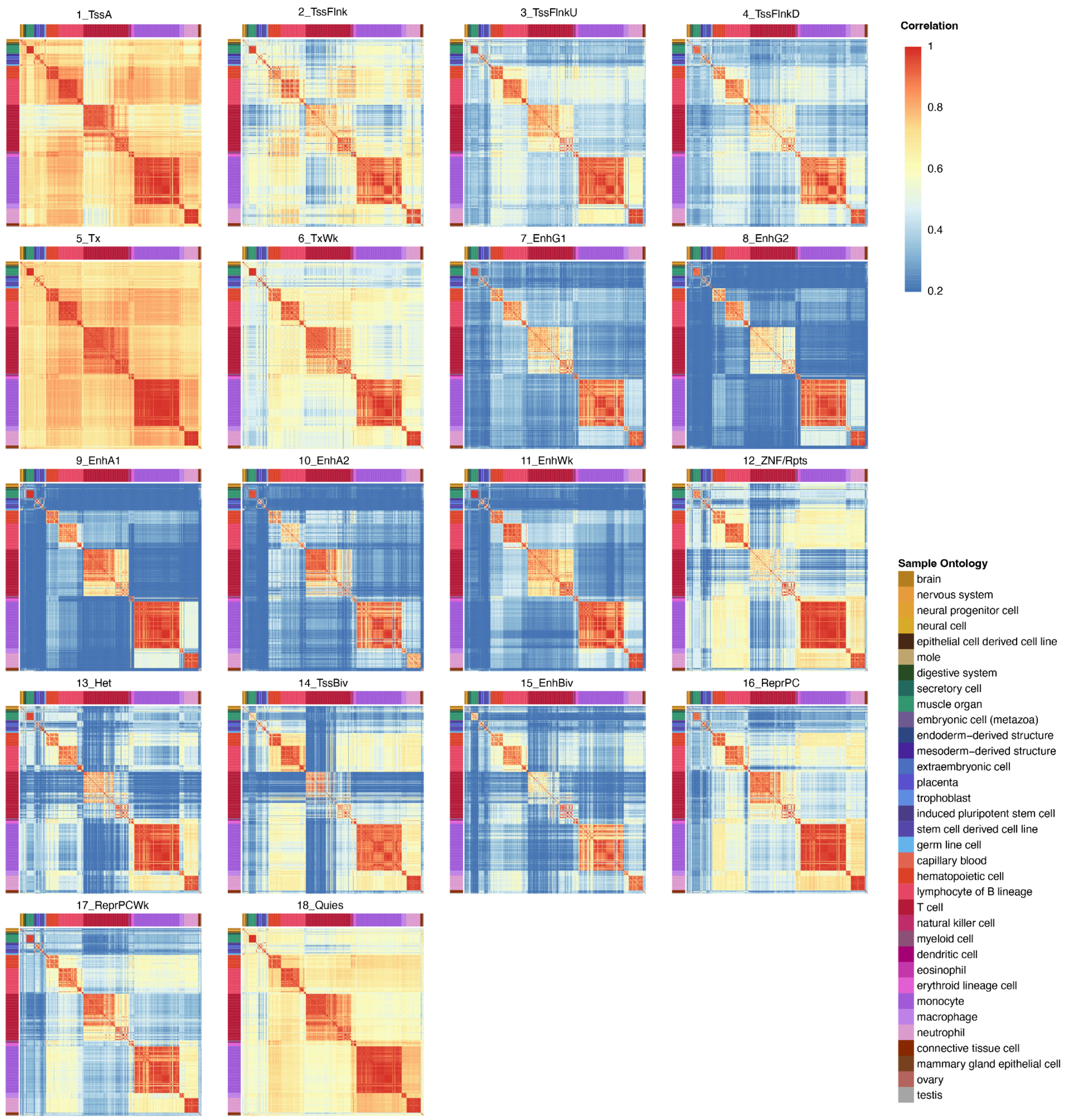


**Supplementary Figure 19: Pairwise sample correlation generated from GECSI-imputed chromatin state soft assignments in 18 chromatin states in the predCS epigenomes.** Heatmaps showing for each of the 18 chromatin states the pairwise correlation calculated using the soft assignments of the chromatin states among 449 GECSI-predicted chromatin states from predCS epigenomes. In each heatmap, each row and column represents an epigenome, and each entry represents a Pearson correlation between the epigenome of the row and the epigenome of the column. The color of the entries follow the color scale on the top right of the figure (0.2–1.0), with values below 0.2 mapped to the color for 0.2. Correlation results are ordered based on the provided sample ordering of IHEC, and color annotation of each sample is based on the sample intermediate ontology in the IHEC metadata using the IHEC coloring.

| State Number and Abbreviation | State Description |
|---|---|
| 1_TssA | Active TSS |
| 2_TssFlnk | Flanking TSS |
| 3_TssFlnkU | Flanking TSS Upstream |
| 4_TssFlnkD | Flanking TSS Downstream |
| 5_Tx | Strong Transcription |
| 6_TxWk | Weak Transcription |
| 7_EnhG1 | Genic Enhancer 1 |
| 8_EnhG2 | Genic Enhancer 2 |
| 9_EnhA1 | Active Enhancer 1 |
| 10_EnhA2 | Active Enhancer 2 |
| 11_EnhWk | Weak Enhancer |
| 12_ZNF/Rpts | ZNF Gene and Repeats |
| 13_Het | Heterochromatin |
| 14_TssBiv | Bivalent/Poised TSS |
| 15_EnhBiv | Bivalent Enhancer |
| 16_ReprPC | Repressed Polycomb |
| 17_ReprPCWk | Weak Repressed Polycomb |
| 18_Quies | Quiescent/Low |

**Supplementary Table 1: Chromatin state description.** Table showing the chromatin state number and abbreviation along with a description based on annotations from the Roadmap Epigenomics Consortium[1].

| Fold | 1 | 2 | 3 | 4 | 5 |
| --- | --- | --- | --- | --- | --- |
| *R* | 10 | 10 | 3 | 10 | 10 |
| *L* | 15 | 10 | 15 | 15 | 15 |
| $\lambda$ | 0.0001 | 0.0001 | 0.0001 | 0.0001 | 0.0001 |

**Supplementary Table 2: Parameter choices for the number of trained models to ensemble (*R*),the number of features for each individual model (*L*), and the lasso penalty parameter ($\lambda$) in cross-validation.** Table reporting the parameter choices for GECSI based on 5-fold cross-validation tuning result, with *R* representing the number of trained models to ensemble, *L* representing the number of features for each individual model, and $\lambda$ representing the lasso penalty parameter.

| Fold | 1 | 2 | 3 | 4 | 5 |
| --- | --- | --- | --- | --- | --- |
| *k* | 7 | 7 | 5 | 7 | 9 |

**Supplementary Table 3: Parameter choices for K-nearest-neighbor (KNN) method.** Table reporting the parameter choices for KNN based on 5-fold cross-validation tuning result, with *k* representing the number of nearest samples to consider.
